# Compound- and fiber type-selective requirement of AMPKγ3 for insulin-independent glucose uptake in skeletal muscle

**DOI:** 10.1101/2021.03.05.433354

**Authors:** Philipp Rhein, Eric M. Desjardins, Ping Rong, Danial Ahwazi, Nicolas Bonhoure, Jens Stolte, Matthieu D. Santos, Ashley J. Ovens, Amy M. Ehrlich, José L. Sanchez Garcia, Qian Ouyang, Mads F. Kjolby, Mathieu Membrez, Niels Jessen, Jonathan S. Oakhill, Jonas T. Treebak, Pascal Maire, John W. Scott, Matthew J. Sanders, Patrick Descombes, Shuai Chen, Gregory R. Steinberg, Kei Sakamoto

**Affiliations:** Nestlé Research, Société des Produits Nestlé S.A., EPFL Innovation Park, Lausanne, 1015, Switzerland; School of Life Sciences, EPFL Innovation Park, Lausanne, 1015, Switzerland; Centre for Metabolism, Obesity, and Diabetes Research, McMaster University, Hamilton, ON, L8N3Z5, Canada; Department of Medicine and Department of Biochemistry and Biomedical Sciences, McMaster University, Hamilton, ON, L8N3Z5, Canada; MOE Key Laboratory of Model Animal for Disease Study, Model Animal Research Center, School of Medicine, Nanjing University, Nanjing, 210061, China; Novo Nordisk Foundation Center for Basic Metabolic Research, University of Copenhagen, Copenhagen, 2200, Denmark; Université de Paris, Institut Cochin, INSERM, CNRS. 75014 Paris, France; Metabolic Signalling Laboratory, St Vincent’s Institute of Medical Research, School of Medicine, University of Melbourne, Fitzroy, VIC 3065, Australia; Mary MacKillop Institute for Health Research, Australian Catholic University, Fitzroy, VIC 3000, Australia; Department of Biomedicine, Aarhus University, Aarhus Denmark, Department of Clinical Pharmacology and Steno Diabetes Center Aarhus, Aarhus University Hospital, Aarhus, Denmark; Protein Chemistry and Metabolism Unit, St Vincent’s Institute of Medical Research, Fitzroy, VIC 3065, Australia; The Florey Institute of Neuroscience and Mental Health, Parkville, VIC 3052, Australia

**Keywords:** AMP-activated protein kinase, 5-aminoimidazole-4-carboxamide riboside, MK-8722, glucose uptake, TBC1D1, Brown adipose tissue

## Abstract

**Objective:** The metabolic master-switch AMP-activated protein kinase (AMPK) mediates insulin-independent glucose uptake in muscle and regulates the metabolic activity of brown and beige adipose tissue (BAT). The regulatory AMPKγ3 isoform is uniquely expressed in skeletal muscle and also potentially in BAT. Here, we investigated the role that AMPKγ3 plays in mediating skeletal muscle glucose uptake and whole-body glucose clearance in response to small-molecule activators that act on AMPK via distinct mechanisms. We also assessed if γ3 plays a role in adipose thermogenesis and browning.

**Methods:** Global AMPKγ3 knockout (KO) mice were generated. A systematic whole-body, tissue and molecular phenotyping linked to glucose homeostasis was performed in γ3 KO and wild type (WT) mice. Glucose uptake in glycolytic and oxidative skeletal muscle *ex vivo,* as well as blood glucose clearance in response to small molecule AMPK activators that target nucleotide-binding domain of γ subunit (AICAR) and allosteric drug and metabolite (ADaM) site located at the interface of the α and β subunit (991, MK-8722) were assessed. Oxygen consumption, thermography, and molecular phenotyping with a β3-adrenergic receptor agonist (CL-316,243) treatment were performed to assess BAT thermogenesis, characteristics and function.

**Results:** Genetic ablation of γ3 did not affect body weight, body composition, physical activity, and parameters associated with glucose homeostasis under chow or high fat diet. γ3 deficiency had no effect on fiber-type composition, mitochondrial content and components, or insulin-stimulated glucose uptake in skeletal muscle. Glycolytic muscles in γ3 KO mice showed a partial loss of AMPKα2 activity, which was associated with reduced levels of AMPKα2 and β2 subunit isoforms. Notably, γ3 deficiency resulted in a selective loss of AICAR-, but not MK-8722-induced blood glucose-lowering *in vivo* and glucose uptake specifically in glycolytic muscle *ex vivo*. We detected γ3 in BAT and found that it preferentially interacts with α2 and β2. We observed no differences in oxygen consumption, thermogenesis, morphology of BAT and inguinal white adipose tissue (iWAT), or markers of BAT activity between WT and γ3 KO mice.

**Conclusions:** These results demonstrate that γ3 plays a key role in mediating AICAR- but not ADaM site binding drug-stimulated blood glucose clearance and glucose uptake specifically in glycolytic skeletal muscle. We also showed that γ3 is dispensable for thermogenesis and browning of iWAT.

## 1. INTRODUCTION

AMP-activated protein kinase (AMPK) is an evolutionary conserved energy sensor that functions to maintain energy homeostasis through coordinating metabolic pathways [1; 2]. AMPK exists as complexes of three subunits; a catalytic α and two regulatory β and γ subunits. Each exists as multiple isoforms (α1/α2, β1/β2, γ1/γ2/γ3), generating up to twelve possible combinations [1]. AMPK heterotrimers are active when a conserved threonine (Thr172) residue within the activation loop of the α subunit kinase domain is phosphorylated [3]. The major upstream kinase phosphorylating Thr172 in metabolic tissues (e.g., muscle, liver) is a complex containing LKB1 [4; 5]. The γ-subunits contain four tandem cystathionine β-synthase (CBS) motifs that bind adenine nucleotides. Binding of ADP and/or AMP to CBS motifs causes conformational changes that promote net Thr172 phosphorylation [6–8]. Moreover, the binding of AMP, but not ADP, further increases AMPK activity by direct allosteric stimulation [6]. Prodrugs of AMP-mimetics such as 5-aminoimidazole-4-carboxamide riboside (AICAR) have been widely used as pharmacological AMPK activators that target the CBS motifs [9]. Proof of concept preclinical studies demonstrated that AICAR treatment improved insulin sensitivity in animal models of insulin resistance [10]. However, AICAR produces numerous AMPK-independent metabolic actions [11]. For example, we have recently demonstrated that AICAR suppresses hepatic glucose production independently of AMPK [12] through inhibition of fructose-1,6-bisphosphatase-1, an AMP-sensitive enzyme involved in gluconeogenesis, *in vivo* [13]. We also showed that AICAR regulated >750 genes in AMPK-null mouse primary hepatocytes [14].

A nucleotide-independent regulation of AMPK was discovered when a novel small-molecule activator, A-769662, was identified [15] and mechanism of action explored [16–18]. The crystallographic structures of AMPK trimeric complexes revealed that A-769662 and 991 (another activator, also known as ex229) bind in a pocket termed allosteric drug and metabolite (ADaM) site located at the interface of the α subunit (kinase domain N-lobe) and β subunit (carbohydrate binding module) [9; 19; 20]. A-769662 was subsequently found to be selective for the AMPKβ1-containing complexes [17] and failed to stimulate AMPK-dependent glucose uptake due to lack of potency against β2-containing complexes that are prevalent in skeletal muscle [21]. We and others have shown that 991, and its two related benzimidazole derivatives with improved bioavailability (MK-8722, PF-739), are potent and highly-specific AMPK activators [14; 22; 23]. They activate both β1- and β2-containing complexes (thereby activating all twelve possible human AMPK complexes) and have been shown to stimulate glucose uptake in skeletal muscle and lower blood glucose levels *in vivo* [22; 24]. Notably, administration of PF-739 resulted in attenuated blood glucose reduction in skeletal muscle-specific but not in liver-specific double knockout of AMPKα1/α2 [23].

AMPK isoform expression varies among different cell and tissue types, with α1, β1, and γ1 appearing the most ubiquitously expressed. Conversely, γ3 is selectively expressed in skeletal muscles containing a high proportion of glycolytic/fast-twitch fibers such as extensor digitorum longus (EDL) muscle [22; 25-27]. Interestingly, even though skeletal muscle expresses multiple isoforms, assays of immunoprecipitated isoforms reveal the α2β2γ1 and α2β2γ3 complexes account for 90% (of which α2β2γ3 accounts for 20%) of the total AMPK trimers in mouse EDL skeletal muscle [21]. Loss of expression/function of α2, β2 or γ3 is sufficient to ablate AICAR-induced glucose uptake in isolated skeletal muscle *ex vivo* [25; 28-32].

In addition to its established metabolic roles in skeletal muscle [33; 34], AMPK also plays a vital role in regulating the development of brown adipose tissue (BAT), maintenance of BAT mitochondrial function, and browning of white adipose tissue (WAT) [35]. Adipose-specific AMPKβ1/β2-null (ad-AMPK KO) mice had a profound defect in thermogenesis [36], and both cold exposure and acute treatment with the β3-adrenergic receptor agonist (CL-316,243) in the ad-AMPK KO mice yielded subnormal increments in oxygen consumption and BAT temperature responses (likely related to impairments in BAT mitochondrial function). A high-throughput screen of protein kinases using a combination of RNAi-mediated knockdown and pharmacological inhibitors identified AMPK as a prominent kinase that promoted the formation of UCP1-abundant brown adipocytes *in vitro* [37]. Proof of concept experiments *in vivo* demonstrate that daily treatment of diabetic ZDF rats with an AMPK activator (C163, for 6 weeks) increased the formation of brown adipocytes [37]. Intriguingly, transcripts of the *Prakg3* (AMPKγ3 gene) were identified in brown adipocyte precursors at intermediate levels, and RNAi-mediated knockdown of *Prakg3* was sufficient to profoundly block the brown adipocyte formation without affecting general adipose differentiation [37]. These results prompted us to assess if γ3 plays a role in adipose thermogenesis and browning *in vivo*.

We hypothesized that γ3-containing complexes play an important role for insulin-independent and AMPK activator-mediated glucose uptake in skeletal muscle and for regulating BAT thermogenesis. To test this hypothesis, we generated γ3 KO mice and determined the effect of AICAR and the ADaM site binding drugs (991, MK-8722) on glucose uptake in glycolytic and oxidative skeletal muscles *ex vivo* and also blood glucose kinetics *in vivo*. In addition, we probed BAT function using the β3-AR agonist CL-316,243. Strikingly, we found that γ3 deficiency resulted in a selective loss of AICAR-, but not 991/MK-8722-induced blood glucose clearance *in vivo* and glucose uptake specifically in glycolytic muscle *ex vivo*. We also found that γ3 is not required for the acute induction of UCP1-mediated non-shivering thermogenesis in the BAT, for the adaptive response to non-shivering thermogenesis or the browning of WAT.

## 2. MATERIALS AND MTHODS

### 2.1. Materials

5-aminoimidazole-4-carboxamide riboside (AICAR) was purchased from Apollo Scientific (OR1170T; Bredbury, United Kingdom). 991 (5-[[6-chloro-5-(1-methylindol-5-yl)-1H-benzimidazol-2-yl]oxy]-2-methyl-benzoic acid) (CAS#: 129739-36-2) was synthesized by Spirochem (Basel, Switzerland) as previously described [1]. Protein G Sepharose and P81 paper were purchased from GE Healthcare (Chicago, IL, USA). [γ-^32^P]-ATP was purchased from PerkinElmer (Waltham, MA, USA). The substrate peptide AMARA was synthesized by GL Biochem (Shanghai, China). All other reagents were from MilliporeSigma (Burlington, MA, USA) if not otherwise stated. Lists of primary and secondary antibodies are in the **Supplementary Table 1 and 2**.

### 2.2. Animal ethics and models

Animal experiments were approved by the internal and local ethics committee and conducted in accordance with the European Convention for the Protection of Vertebrate Animals used for Experimental and Other Scientific Purposes. Protocols used were approved by the Service Vétérinaire Cantonal (Lausanne, Switzerland) under licenses VD3332 and VD3465, and also approved by an ethical committee (Com’Eth, CE17) registered at the French Ministry of Research under the reference: 10261, as well as were in accordance with McMaster Animal Care Committee guidelines (AUP #: 16-12-41, Hamilton, ON), and the Ethics Committees at Nanjing University with involved personnel having personal licenses from the regional authority. The generation of a constitutive *Prkag3*^−/−^ (AMPKγ3^−/−^) mice was performed by Taconic Biosciences as described in **Supplementary Figure 1**. The AMPKα1f/f and AMPKα2f/f mice were as previously described [38], and obtained from the Jackson Laboratory (Bar Harbor, ME, USA). These two strains were used to derive AMPKα1f/f/α2f/f mice that were then bred with the Mlc1f-Cre mice to obtain the AMPKα1f/f/α2f/f - Mlc1f-Cre mice. The resultant AMPKα1f/f/α2f/f - Mlc1f-Cre mice are the AMPKα1/α2 skeletal muscle-specific knockout mice. TBC1D1 S231A knock-in mice have been described [39]. All these lines are on C57BL6 background. The animals were kept and maintained according to local regulations under a light-dark cycle of 12 hours and had free access to a standard chow diet. Male mice ranging 10-16 weeks of age were used for experiments otherwise stated. High fat diet (60% kcal% fat) was obtained from Research Diet (RD 12492).

### 2.3. Analysis of body composition and plasma hormone levels

Body composition (fat content, lean tissues and free body fluid) was assessed using the Minispec analyser (Bruker) by Nuclear Magnetic Resonance (NMR) technology. The test was conducted on conscious fed mice. Blood was collected at the indicated age by retro orbital puncture under isoflurane anesthesia at noon on mice fasted for 4 hours. Plasma insulin and leptin levels were measured on a BioPlex analyser (BioRad) using the Mouse Metabolic Magnetic Hormone Magnetic Bead panel kit (MilliporeSigma).

### 2.4. Oral glucose tolerance test

Mice were fasted overnight (16 hours) and a bolus of glucose solution (2 g/kg body weight) was administered via oral gavage. Blood glucose collected from tail vein was measured at different time points over 120 min using blood glucose monitor and glucose test strips (Roche Diagnostics, Accu-Chek).

### 2.5. Preparation of mouse tissue extracts for protein analysis

Mouse tissue were dissected and immediately frozen in liquid nitrogen. The tissues were homogenized in ice-cold lysis buffer (270 mM sucrose, 50 mM Tris·HCl (pH 7.5), 1 mM EDTA, 1 mM EGTA, 1% (w/v) Triton X-100, 20 mM glycerol-2-phosphate, 50 mM NaF, 5 mM Na_4_P_2_O_7_, 1 mM DTT, 0.1 mM PMSF, 1 mM benzamidine, 1 μg/ml microcystin-LR, 2 μg/ml leupeptin, and 2 μg/ml pepstatin A) using a tissue lyser (Tissue Lyser II; Qiagen). Lysates were centrifuged at 21,300 *g* for 15 min and protein concentration from the supernatant was determined using Bradford reagent (23200, ThermoFisher) and bovine serum albumin (BSA) as standard. The supernatants were stored in aliquots in −80°C freezer until subsequent analysis.

### 2.6. Immunoblotting

Protein extracts were denatured in Laemmli buffer at 95°C for 5 min. 20 μg of protein was separated by SDS-PAGE on 4-12% gradient gels (NW04127, ThermoFisher) and transferred onto nitrocellulose membranes (#926-31090, LiCOR). Membranes were blocked for 1 hour at room temperature in LiCOR blocking buffer (#927-60001, LiCOR). The membranes were subsequently incubated in TBST (10 mM Tris (pH 7.6), 137 mM NaCl, and 0.1% (v/v) Tween-20) containing 5% (w/v) BSA and the primary antibody overnight at 4°C. After extensive washing, the membranes were incubated for 1 hour in either HRP-conjugated or LiCOR secondary antibodies diluted 1:10,000. Signal imaging was performed either using enhanced chemiluminescence (ECL) reagent (GE Healthcare) or a LiCOR Odyssey CLx imaging system. Densitometry for ECL blots was performed using Image J Software (NIH). Due to sample limitation (from the incubated muscle tissue samples), we also utilized automated capillary Western Blot system Sally Sue (ProteinSimple, San Jose, CA, USA). Experiments were performed according to the manufacturer’s protocol using the indicated standard reagents for the Sally Sue system (SM-S001, ProteinSimple). Briefly, all samples were first diluted to 2 mg/ml in lysis buffer and then further diluted to 0.5 mg/ml in 0.1% SDS. Following the manufacturer’s instructions for sample preparation, this resulted in an assay protein concentration of 0.4 mg/ml.

### 2.7. Immunoprecipitation and in vitro AMPK activity assay

Lysates of muscle (200 μg) or BAT (1,000 μg) were incubated on a rotating platform at 4°C overnight with a mix of 5 μl protein G-sepharose and the indicated antibodies. The beads were pelleted at 500 *g* for 1 min and initially washed twice with 0.5 mL lysis buffer containing 150 mM NaCl and 1 mM DTT and subsequently washed twice with the same amount of buffer A [50 mM HEPES (pH 7.4), 150 mM NaCl, 1 mM EGTA and 1 mM DTT]. The AMPK complexes were either eluted with Laemmli buffer for immunoblot analysis or taken directly for AMPK activity measurement. The AMPK activity assay was performed by incubating the beads (immune-complexes) for 30 min at 30°C on a heated shaker in buffer A with additional 10 mM Mg^2+^ and 100 μM ATP in presence of 200 μM AMARA peptide (AMARAASAAALARRR) and 1 μCi of [γ-^32^P] ATP [22]. Reactions were stopped by spotting the reaction mix onto P81 filter paper and washing in 75 mM phosphoric acid. The P81 papers were dried after three washes and the ^32^P incorporation into the substrate peptide measured by Cherenkov counting (5 min) using a scintillation counter (Tri-Carb 2810TR, PerkinElmer).

### 2.8. Citrate synthase activity assay

Protein extracts (10 µg) (using the same lysis buffer described above) was assayed in duplicates using a citrate synthase assay kit (CS0720, MilliporeSigma) according to the manufacturer’s instruction using recombinant citrate synthase as positive control.

### 2.9. Analysis of gene expression and mitochondrial DNA copy number using quantitative real-time PCR (qPCR)

To perform a relative quantification of mRNA levels of the AMPK subunit isoforms in mouse skeletal muscle tissues, reverse transcription and qPCR was performed as described [14]. All the primers and sequences are listed in the **Supplementary Table 3**. Relative mRNA quantities were calculated for triplicate muscle samples from 4-5 animals and normalized using the three reference genes *Hprt1* (hypoxanthine ribosyltransferase, HPRT), *GusB* (beta-glucuronidase) and *Pgk1* (Phosphoglycerate Kinase 1). Real-time qPCR in brown adipose tissue was performed separately as described [36]. Relative gene expression was calculated using the comparative Ct (2-ΔCt) method, where values were normalized to a reference gene (*Ppia*).

To relatively quantify the amount of mtDNA present per nuclear genome by qPCR, mtDNA (16S, ND4) and nuclear DNA (PMP22, Titin) primers and probes were used, the sequences of which are shown in **Supplementary Table 3**. The relative mt copy number was determined based on the relative abundance of nuclear and mtDNA, calculated as average of the two targets respectively. The relative abundance is then expressed by ΔCT or CT(nDNA) - CT(mtDNA) and displayed as fold change of copy number of mtDNA per nuclear genome compared to the WT muscle.

### 2.10. Immunofluorescence for fiber type determination and fiber size

For immunostaining against Myh4, Myh2, Myh7 and Laminin, mouse hindlimbs (no skin) without fixation were embedded with Tissue-TEK OCT (Sakura Finetek, Netherlands) and directly frozen in cold isopentane pre-cooled in liquid nitrogen as described [40]. Hindlimb cross sections were prepares using a cryostat (Leica 3050s) with a thickness of 10 μm. The cross sections were washed 3 times for 5 min with PBS and then incubated with blocking solution (PBS and 10% goat serum) for 30 min at room temperature. Sections were incubated overnight with primary antibody diluted in PBS + 10% goat serum solution at 4°C and washed as described above. The sections were incubated with secondary antibody, diluted in PBS + 10% goat serum solution for 1 hour at room temperature. Sections were further washed and mounted with Mowiol solution and a glass coverslip. Images were collected with a microscope (Olympus BX63F) and a camera (Hamamatsu ORCA-Flash 4.0). Images were analyzed with ImageJ (NIH). Fiber boundaries were defined by the laminin signal and myosin Myh7 (type I), Myh2 (type IIA) and Myh4 (type IIB) heavy chains were quantified. Remaining unlabeled fibers were included for total fiber number and individual proportions of type I, type IIA and type IIB of that total number calculated.

### 2.11. Ex vivo skeletal muscle incubation and analysis of glucose uptake

Animals were anesthetized by Avertin [2,2,2-Tribromoehtanol (Sigma-Aldrich #T48402) and 2-Methyl-2-butanol 99% (Sigma-Aldrich #152463)] via intraperitoneal injection, and EDL or soleus muscles were rapidly dissected and mounted in oxygenated (95% O_2_ and 5% CO_2_), and warmed (30°C) Krebs-Ringer buffer in a myograph system (820MS DMT, Denmark). The respective muscles were incubated as described [22; 41] in the presence of the indicated drug or vehicle for 50 min. 2-deoxy-[^3^H] glucose uptake was measured during the last 10 min of the incubation as described [22; 39].

### 2.12. AICAR and MK-8722 tolerance test

Access of the mice to food was restricted for 3 hours (07:00-10:00) prior to the experiment. AICAR (250 mg/kg body weight) or vehicle (water) was injected intraperitoneally and blood glucose levels were monitored for 120 min using the Contour XT glucometer (Bayer, Leverkusen) and single use glucose sensors (Ascensia, Basel). MK-8722 tolerance was tested by oral administration of either MK-8722 (10 or 30 mg/kg body weight) or vehicle (0.25% (w/v) methylcellulose, 5% (v/v) Polysorbate 80, and 0.02% (w/v) sodium lauryl sulfate in deionized water) [24]. Blood glucose measurement was performed as described above for AICAR.

### 2.13. ZMP and adenine nucleotide measurements

Muscle tissues were lysed in 200 μl cold 0.5 M perchloric acid. Extracts were collected and clarified at 14,000 rpm for 3 min. 100 μl clarified lysate was neutralized with 25 μl cold 2.3 M KHCO_3_ and incubated on ice for 5 min. Samples were centrifuged at 14,000 rpm for 3 min. ZMP were measured by LC-MS/MS with modifications to our previously described method [42]. Both LC and MS instruments were controlled and managed with the Analyst 1.7.1 software (AB Sciex). The autosampler was set at 4°C and column oven set at 30°C, which housed a 150 mm (length) x 0.5 mm (inner diameter) Hypercarb 3 μm porous graphitic carbon column (Thermo Fisher Scientific). The LC solvent system comprised of 50 mM triethylammonium bicarbonate buffer (TEAB, MilliporeSigma) pH 8.5 in pump A, and acetonitrile with 0.5 % trifluoroacetic acid (TFA; Sigma-Aldrich) in pump B. A flow rate of 400 μl/min was used throughout a gradient program consisting of 0 % B (2 min), 0 to 100 % B (10 min), 100 % B (3 min), 0 % B (2 min). Data was analyzed with MultiQuant 3.0.2 software (AB Sciex) using area under the LC curve. Calibration curves were determined by linear regression of the peak area of a ZMP standard curve and were required to have a correlation coefficient (R^2^) of > 0.98. Adenine nucleotides and adenylate energy charge were measured by LC-MS as described [43].

### 2.14. Adipose tissue histology

Tissues were fixed in 10% formalin for 24-48 hours at 4°C and processed for paraffin embedding and hematoxylin and eosin staining by the core histology laboratory at the McMaster Immunology Research Centre (Hamilton, Canada).

### 2.15. Infrared thermography

UCP1-mediated thermogenesis was assessed in 14-week-old male wild-type and γ3 KO mice as described [44]. Briefly, mice were anaesthetized with an intraperitoneal injection of 0.5 mg/g body weight Avertin (2,2,2-Tribromoethanol dissolved in 2-methyl-2-butanol; MilliporeSigma) then, 2 min later, injected with either saline or the highly selective β3-adrenergic receptor agonist CL 316,243 (Tocris, Bristol, United Kingdom). Mice were subsequently placed dorsal side up onto an enclosed stationary treadmill to measure oxygen consumption (VO_2_) with a Comprehensive Laboratory Animal Monitoring System (Columbus Instruments, OH, USA) and 18 min after the second injection a static dorsal thermal image was taken with an infrared camera (FLiR Systems, Wilsonville, OR, USA). Serum samples were collected via tail-nick just after the infrared image was taken and non-esterified free fatty acid (NEFA) concentration was determined using manufacturer instructions with a two-step kit (NEFA-HR 2, WAKO).

### 2.16. Metabolic monitoring

Metabolic monitoring was performed as described [45] in a Comprehensive Laboratory Animal Monitoring System. For the chronic 5-day CL 316,243 challenge, mice were injected intraperitoneally with saline or CL 316,243 (first 4 days 0.5 mg/kg and last day 1.0 mg/kg) at 09:30 hours and measurements for VO_2_ were calculated 6 hours post-injection. Mice were euthanized 24 hours after the last injection.

### 2.17. Statistical analysis

Data are reported as mean ± SEM and statistical analysis was performed using GraphPad Prism software. As indicated in the respective figure legends, differences between only two groups were analyzed using an unpaired two-tailed Student’s *t*-test and for multiple comparisons one-way analysis of variance (ANOVA) with Bonferroni *post hoc* test or repeated measures two-way ANOVA was used. Statistical significance was accepted at *P* < 0.05.

## 3. RESULTS

### 3.1. A genetic/constitutive loss of the AMPKγ3 reduced AMPKα2 and β2 protein abundance in mouse glycolytic skeletal muscle

We generated constitutive AMPKγ3 KO mice through flanking exons 5-10 of *Prkag3* gene with LoxP sites (**Supplementary Fig. 1A**). This is expected to cause a loss-of-function of the *Prkag3* gene by deleting the nucleotide binding cystathionine β-synthase (CBS)-2 domain and parts of the CBS-1 and -3 domains and by generating a frame shift from exon 4 to exon 11 (premature stop codon in exon 12). In addition, the resulting transcript may be a target for non-sense mediated RNA decay and thereby may not be expressed at significant level. In support of this, we were unable to detect faster migrating polypeptides using the antibody raised against residues 44–64 (within exon 1-3) of the mouse γ3 (**Supplementary Fig. 1B**). γ3 homozygous KO (γ3^−/−^) mice were born at expected Mendelian frequency (data not shown). Food intake and spontaneous physical activity, as well as oxygen consumption were similar between wild-type (WT) and γ3 KO mice (**Supplementary Fig. 1C-E**).

We first confirmed a complete loss of γ3 protein and its associated AMPK catalytic activity in tissues harvested from γ3 KO mice (**Fig. 1A, B**). Expression of γ3 is restricted to skeletal muscles containing a high proportion of glycolytic/fast-twitch fibers [22; 25]. In line with this, we observed that γ3 and its associated AMPK activity were predominantly detected in glycolytic gastrocnemius (GAS) and extensor digitorum longus (EDL) muscles in WT mice. A modest expression of γ3 and its associated AMPK activity were detected in soleus muscle (which contains a high proportion of oxidative/slow-twitch fibers) from WT mice when γ3 proteins were enriched by immunoprecipitation prior to the immunoblotting (**Fig. 1A, B**). We detected a faint band immuno-reactive to the γ3 antibody in liver lysates from both WT and γ3 KO mice (**Fig. 1A**, middle panel). We confirmed that the observed band was non-specific as it was readily detected in IgG control samples (**Fig. 1A**, lower panel) and only a negligible background γ3-associated AMPK activity was detected in liver (and also heart) lysates in both WT and γ3 KO mice (**Fig. 1B**). Next, we assessed if loss of γ3 affected AMPKα1- and α2-containing complex activity in GAS muscle. As illustrated in **Fig 1C and D**, we observed that while AMPKα1 activity was unaltered, AMPKα2 activity was reduced (∼50%). Since we and others have shown that γ3 predominantly interacts with α2 and β2 [21; 22] to form a stable trimeric α2β2γ3 complex, we hypothesized that a constitutive loss of γ3 would cause reduced expressions of α2 and β2 due to their destabilization as monomers. To test this hypothesis, we performed an analysis of AMPK subunit/isoform abundance in both glycolytic (EDL and GAS) and oxidative (soleus) muscles in WT and γ3 KO mice (**Fig. 1E-G**). We previously performed an extensive antibody validation for all AMPKαβγ isoforms using individual isoform-specific KO mouse tissues as negative controls and also reported that γ2 proteins (UniProt ID: Q91WG5 isoform A) were not detectable in mouse skeletal muscles [22]. Immunoblot analysis revealed that protein levels of α2, total AMPKα using a pan α1/α2 antibody, and β2 isoforms were selectively reduced (∼20-30%) in EDL and GAS (**Fig. 1E, F)**, but not in soleus (**Fig. 1G)**, of γ3 KO as compared to WT mice. There was no compensatory increase in γ1 isoform in γ3 KO muscles. To examine if the reduced protein abundance of α2 and β2 was due to decreased mRNA expression of the *Prkaa2* and *Prkab2* (the genes encoding AMPKα2 and β2, respectively) in the γ3 KO mice, we performed qPCR analyses (**Fig. 1H, I**). We confirmed that *Prkag3* mRNA expression was undetectable in skeletal muscle from γ3 KO mice, and observed that there were no differences in mRNA expressions of other AMPK subunit/isoforms in GAS (**Fig. 1H**) or soleus (**Fig. 1I**) between WT and γ3 KO mice. Taken together, we have demonstrated that a genetic/constitutive loss of AMPKγ3 causes reductions of AMPKα2 and β2 proteins without affecting their mRNA expressions in glycolytic skeletal muscle.

**Figure 1:**
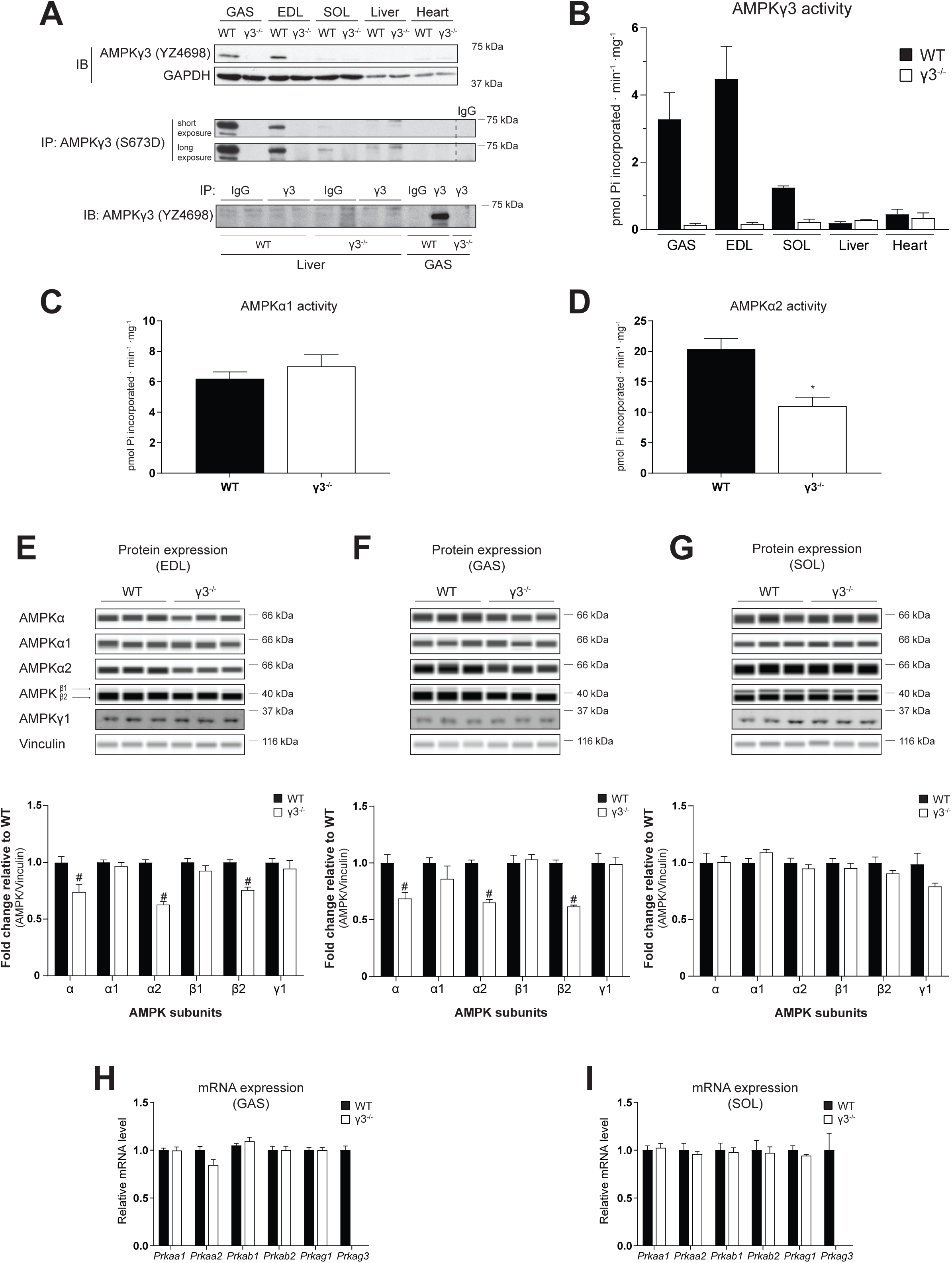
Genetic ablation of the AMPKγ3 causes a significant loss of α2 and β2 expression in mouse glycolytic skeletal muscles. (A) Immunoblot (IB) analysis of γ3 expression in a panel of tissues extracted from wild-type (WT) or AMPKγ3-null (γ3^−/−^) mice (upper panel). γ3 expression was further analyzed by immunoblotting following enrichment of the γ3 proteins via immunoprecipitation (IP) from the indicated tissue extracts (200 μg) (middle panel). Liver and skeletal muscle (GAS) tissue extracts from the indicated genotypes were used for immunoprecipitation with either γ3-specific antibody or species-matched IgG (as negative control) and the immune-complexes were subsequently immunoblotted with γ3 antibody (lower panel). (B) The γ3-containing AMPK complexes were immunoprecipitated from the indicated tissues harvested from the indicated genotypes and an *in vitro* AMPK activity assay was performed in duplicate (n=3 per tissue/genotype). (C, D) The *In vitro* AMPK activity assay was performed on α1- or α2-contaning AMPK complexes immunoprecipitated from GAS extracts (n=9-10 per tissue/genotype). (E-G) Representative immunoblot images and quantification of the AMPK isoform-specific expression using an automated capillary immunoblotting system (Sally Sue) with the indicated antibodies as described in Materials and Methods. AMPK isoform expressions were normalized by their respective vinculin expression (loading control) and are shown as fold change relative to WT. Note that AMPKγ1 expression was quantified using another immunoblotting system (Li-COR, described in the Materials and Methods) due to antibody compatibility (n=5-11 per tissue/genotype). (H, I) Relative levels of mRNA of the indicated genes (encoding AMPK isoforms) in the indicated skeletal muscles were assessed by qPCR (n=5 per tissue/genotype). Results are shown as means ± SEM. Statistical significance was determined using the unpaired, two-tailed Student’s t-test and are shown as ^#^*P* < 0.05 (WT vs. γ3^−/−^). GAS; gastrocnemius, EDL; extensor digitorum longus, SOL; soleus, IgG; immunoglobulin G.

### 3.2. AMPKγ3 deficiency has no impact on mitochondrial content and components or fiber-type composition in skeletal muscle

A loss-of-function of skeletal muscle AMPK is associated with reduced mitochondrial content and function [45–48]. Interestingly, a transgenic mouse model overexpressing γ3 mutant (R225Q, a gain-of-function mutation), was associated with higher mitochondrial content and increased amount of a marker of the oxidative capacity (succinate dehydrogenase) in individual muscle fibers of the white portion of GAS [49]. Nevertheless, γ3 deficiency did not cause alterations in mitochondrial content or other parameters in GAS muscle [49]. However, the previously generated γ3 deficient mice did not exhibit significantly reduced expression or activity of AMPKα2 [25], the predominant α-catalytic isoform in skeletal muscle. In the current study we wanted to address whether γ3 deficiency, coupled to a partial loss of AMPKα2 activity (**Fig. 1D)**, had an impact on mitochondrial parameters in both glycolytic (EDL) and oxidative (soleus) skeletal muscle. We observed no differences in mitochondrial DNA copy number (**Fig. 2 A, B**), citrate synthase activity (**Fig. 2 C, D**), or components of the mitochondrial respiratory chain complex (**Fig. 2E, F**) in soleus or EDL muscles of WT and γ3 mice. Fiber-type analysis of hindlimb cross sections using immunofluorescence revealed no differences in myosin heavy chain isoform composition in EDL or soleus muscles between the genotypes (**Fig. 2G, H**). We also observed no difference in skeletal muscle fiber size (cross sectional area) between the genotypes (**Fig. 2I**). Collectively, we show that constitutive γ3 deficiency does not affect mitochondrial content, respiratory chain complex expression, or fiber type composition in both glycolytic and oxidative skeletal muscle.

**Figure 2:**
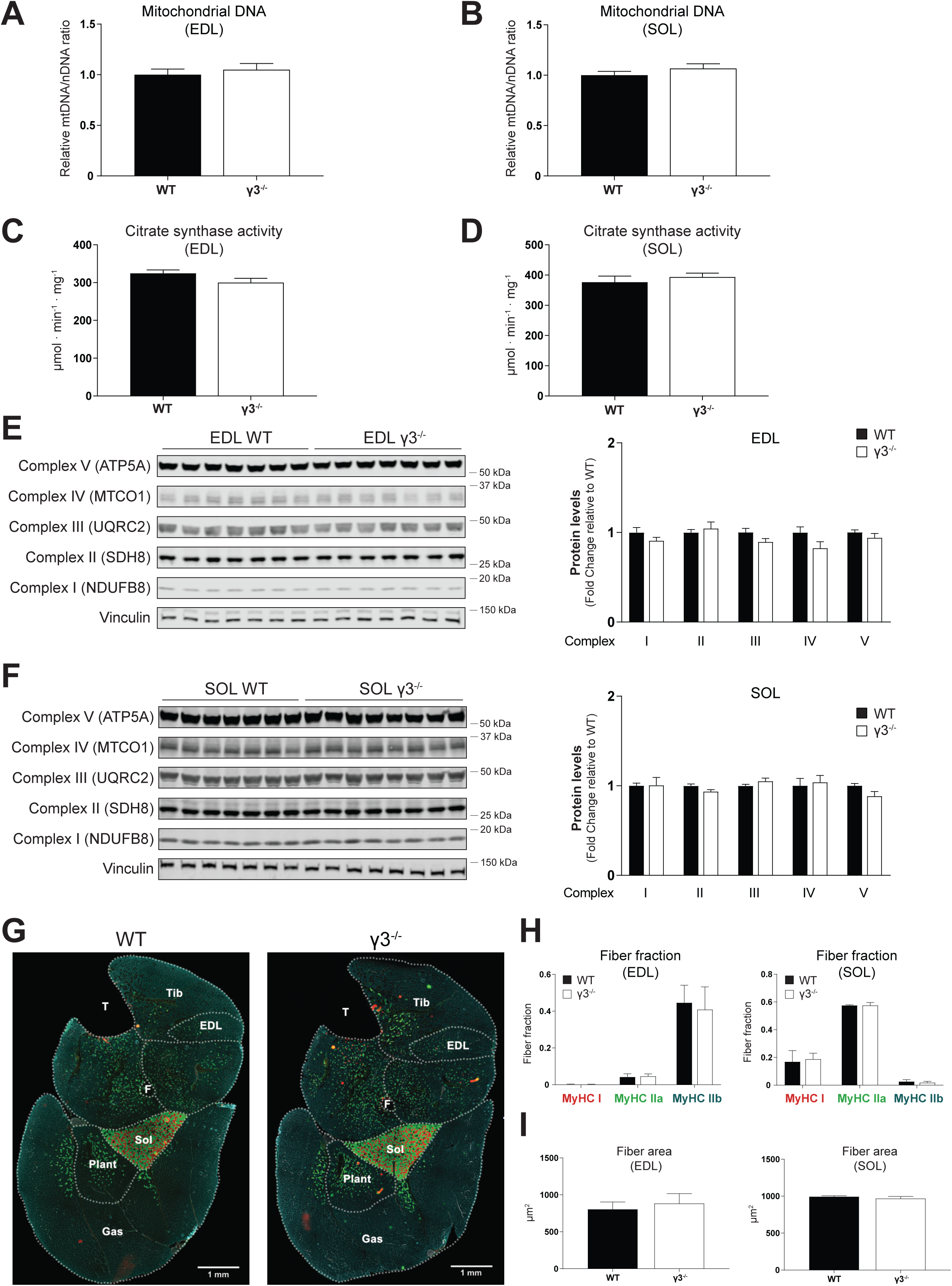
AMPKγ3 deficiency does not affect mitochondrial content and components, or fiber-type composition in skeletal muscles. (A, B) Relative quantification of mitochondrial DNA (mtDNA) was performed using qPCR-based assay as described in the Materials and Methods (n=5 per tissue/genotype). (C, D) Citrate synthase activity was measured in the indicated muscle extracts (n=8 per tissue/genotype). (E, F) Immunoblot analysis and quantification of mitochondrial complexes in the indicated muscles (n=7 per tissue/genotype). (G-I) Representative cross-sectional images (of n=4 per genotype) of the whole-hindlimb muscle fiber-type analysis of the indicated genotypes using isoform-specific myosin heavy chain (MyHC) and laminin antibodies followed by immunofluorescent signal detection (G). Scale bar=1 mm. Quantification of relative isoform-specific myosin heavy chain (MyHC) composition/fraction (red: MyHC I, green: MyHC IIa, blue: MyHC IIb, laminin: gray/white) and fiber area in the indicated muscles were performed as described in Materials and Methods. Unstained fibers are not included in the fiber fraction analysis (H, L, n=3-4 per tissue/genotype). Results are shown as means ± SEM. GAS; gastrocnemius, EDL; extensor digitorum longus; SOL; soleus, Tib; tibialis anterior, Plant; plantaris, F; fibula, T; tibia.

### 3.3. AMPKγ3 KO mice exhibit normal glucose homeostasis on chow and in response to high fat diet (HFD) feeding

Transgenic mice overexpressing the γ3 mutant (R225Q) exhibit an increase in muscle lipid oxidation and are protected against HFD-induced insulin resistance in skeletal muscle [25]. In the current study, we examined if γ3 deficiency affected glucose homeostasis under standard chow and in response to HFD feeding. Body weight and composition were similar between WT and γ3 KO mice during both chow and HFD feeding periods (**Fig. 3A, B**). We observed similar levels of plasma insulin and leptin on chow diet between WT and γ3 KO mice, with their levels increased in a comparable manner for both genotypes in response to HFD feeding (**Fig. 3C, D**). Consistent with these results, we observed comparable fasted blood glucose levels and no difference in glucose tolerance between the genotypes irrespective of the diets (**Fig. 3E**). To complement these *in vivo* results, we assessed insulin signaling and glucose uptake in isolated EDL muscle *ex vivo*. As shown in **Fig. 3F**, basal glucose uptake and Akt phosphorylation were comparable between WT and γ3 KO mice and insulin equally stimulated both parameters in both genotypes. We also confirmed that there was no difference in the expression of GLUT4 and hexokinase II in EDL muscle between WT and γ3 KO mice (**Fig. 3F**). Taken together, these results suggest that γ3 is dispensable for maintenance of glucose homeostasis on chow and in response to HFD.

**Figure 3:**
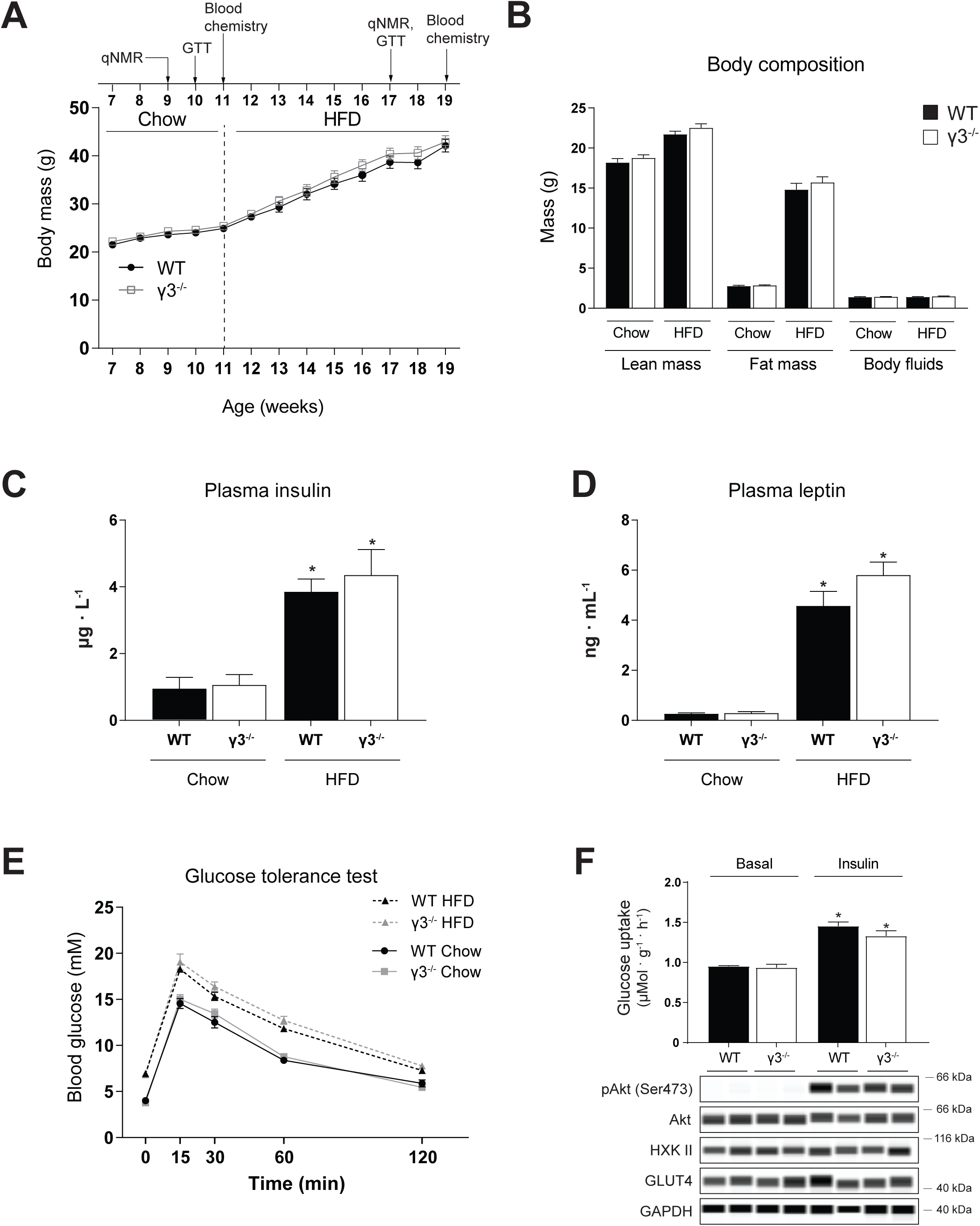
AMPKγ3 is dispensable for maintaining glucose homeostasis under chow and high fat diet (HFD) feeding. (A) Time sequence of the diet intervention, analysis of body composition (qNMR), oral glucose tolerance test (GTT) and plasma hormone analysis (blood chemistry). Mice were fed chow diet after weaning until 11 weeks of age before switching to HFD (60% kcal% fat). Body weight over time of the indicated genotypes (n=10 per genotype). (B) Body composition determined by qNMR in the indicated genotypes during the indicated diet treatment. (C, D) Plasma insulin and leptin levels were determined using the commercial enzyme-linked immunosorbent assay kits. (E) Mice were fasted overnight and an oral GTT test was performed during chow (week 10) and HFD (week 17) feeding by monitoring blood glucose kinetics over the indicated duration following an oral administration of a bolus of glucose solution (2 g/kg body weight). (F) Extensor digitorum longus (EDL) muscles from the indicated genotypes on chow diet (10-12-week old males from a separate cohort. n=5-7 per genotype) were isolated in incubated in the presence or absence of insulin (100 nM) for 50 min and were subjected to glucose uptake assay and immunoblot analysis using the indicated antibodies. Results are shown as means ± SEM. Statistical significance was determined using the unpaired/two-tailed Student’s t-test or one-way analysis of variance with Bonferroni correction and are shown as \**P* < 0.05 (treatment effect within the same genotype).

### 3.4. AMPKγ3 deficiency causes attenuated AICAR-stimulated glucose uptake in glycolytic skeletal muscle ex vivo and blood glucose lowering in vivo

AICAR-stimulated glucose uptake in skeletal muscle requires functional AMPK [34]. Consistent with previous studies [30; 39], AICAR promoted glucose uptake robustly in EDL (∼2.5-fold) and modestly in soleus (∼1.6-fold) *ex vivo* in WT mice (**Fig. 4A, B**). Interestingly, we observed that AICAR-stimulated glucose uptake was profoundly reduced in EDL, but not in soleus, in γ3 KO mice (**Fig. 4A, B**). To examine if a loss of γ3 affected AICAR-induced AMPK activity, we measured phosphorylation of ACC and TBC1D1, established surrogate markers of *cellular* AMPK activity in muscle. As shown in **Fig. 4C-F**, phosphorylation of ACC and TBC1D1 was increased in both EDL and soleus in response to AICAR in WT mice. The AICAR-mediated increase in phosphorylation of ACC and TBC1D1 was reduced in EDL, but not in soleus, in γ3 KO mice (**Fig. 4C-F**). We confirmed that there is no sex-dependent AICAR effect, as AICAR-stimulated glucose uptake was similarly reduced in EDL muscle from female γ3 KO mice (data not shown). We wanted to explore if a partial loss of γ3 results in a reduction of AICAR-stimulated glucose uptake in EDL. Heterozygous γ3^+/−^ mice had ∼50% reduction in γ3 expression in GAS, but the expression of total AMPKα, α2 and β1/β2 was not reduced (**Supplementary Fig. 2A, B**). Incubation of EDL with AICAR *ex vivo* resulted in similar increases in γ3-associated activity and glucose uptake, as well as phosphorylation of ACC and TBC1D1 in both WT and heterozygous γ3^+/−^ mice (**Supplementary Fig. 2C-G**).

**Figure 4:**
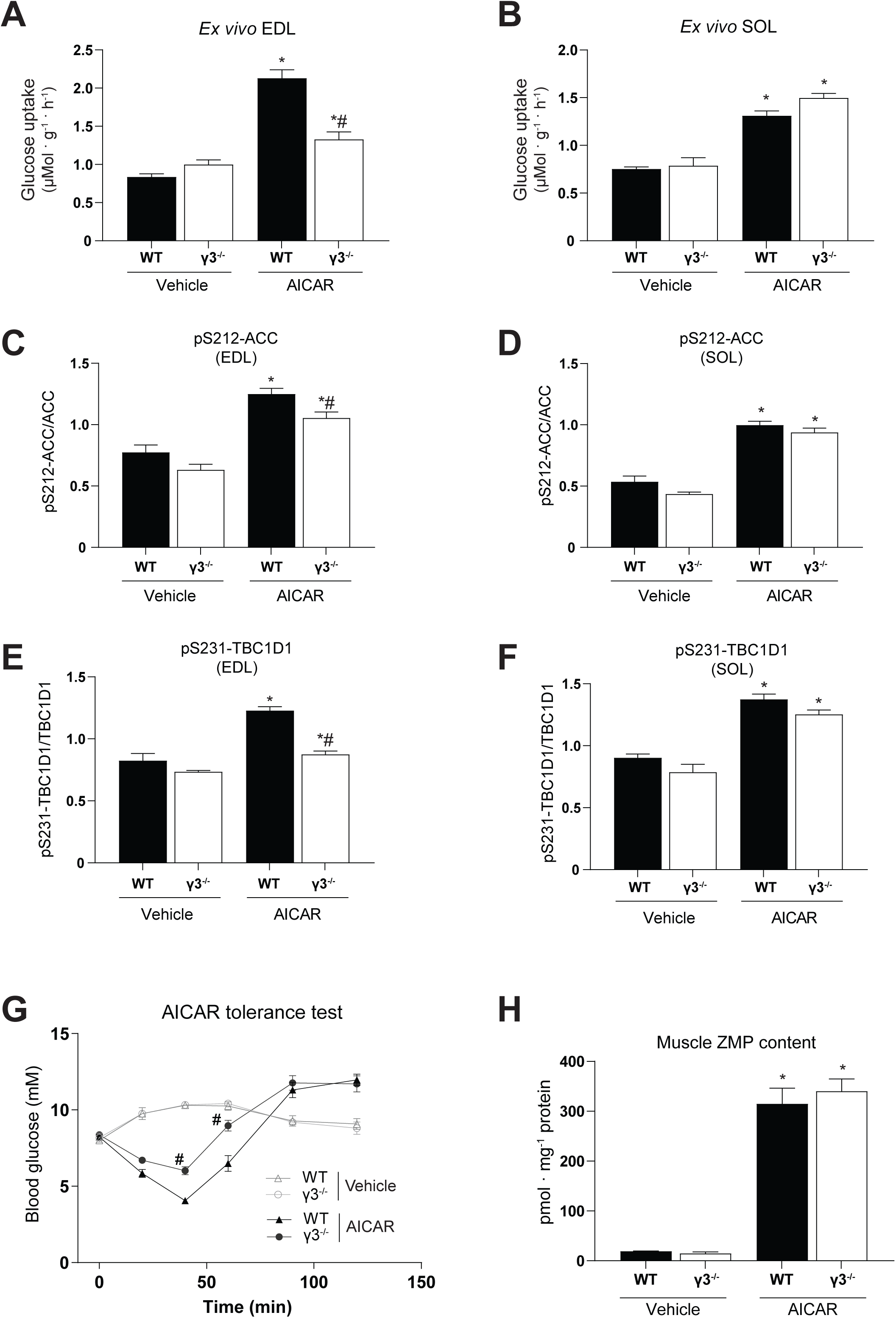
AMPKγ3 is required for AICAR-induced glucose uptake in glycolytic skeletal muscles and hypoglycemia. (A-F) EDL or SOL muscles were isolated from the indicated genotypes and incubated in the absence (vehicle, DMSO) or presence of AICAR (2 mM) for 50 min followed by an additional 10-min incubation with the radioactive 2-deoxy-glucose tracer. One portion of the muscle extracts was subjected to glucose uptake measurement (A, B) and the other was used for immunoblot analysis using the automated capillary immunoblotting system with the indicated antibodies (C-F) (n=4-7 per treatment/genotype). (G, H) AICAR tolerance test and muscle ZMP analysis. Mice were fasted for 3 hours and injected either with vehicle (water) or AICAR (250 mg/kg body weight, i.p.) followed by blood glucose kinetics measurement over the indicated duration (G). Following the AICAR tolerance test, mice were euthanized and GAS muscles were extracted and ZMP levels were determined (H) (n=5-12 per treatment/genotype). Results are shown as means ± SEM. Statistical significance was determined using the unpaired/two-tailed Student’s t-test or one-way analysis of variance with Bonferroni correction and are shown as \**P* < 0.05 (treatment effect within the same genotype), ^#^*P* < 0.05 (WT vs. γ3^−/−^ within the same treatment). GAS; gastrocnemius, EDL; extensor digitorum longus, SOL; soleus, AICAR; 5-aminoimidazole-4-carboxamide ribonucleoside, ZMP; AICAR monophosphate

We next wanted to determine if γ3 deficiency affected the hypoglycemic effects of AICAR *in vivo*. We utilized partially fasted animals (3-hour fast, 07:00-10:00), as the AICAR-induced reduction of blood glucose in overnight fasted (16 hours) mice was predominantly caused by the suppression of hepatic glucose output [13]. After administration of a bolus of AICAR (250 mg/kg body weight, i.p.) or vehicle, we monitored blood glucose kinetics for 2 hours in WT and γ3 KO mice. As shown in **Fig. 4G**, we observed that the blood glucose-lowering action of AICAR was blunted (40 and 60 min time points) in γ3 KO compared to WT mice. We confirmed that ZMP content in GAS muscle following AICAR administration was comparably increased between the two genotypes (**Fig. 4H**). Additionally, AICAR did not affect adenylate energy charge in GAS from both genotypes (**Supplementary Fig. 2H**). Collectively, these results suggest that γ3 (i.e. γ3-containing AMPK complex(es)) plays an important role in AICAR-mediated glucose uptake and disposal in glycolytic skeletal muscles.

### 3.5. ADaM site-targeted activators normally stimulate glucose uptake in skeletal muscle and lower blood glucose levels in AMPKγ3 KO mice

The ADaM site-binding pan AMPK activator, 991, robustly stimulates glucose uptake in isolated mouse skeletal muscle tissues *ex vivo* [22; 50]. We initially confirmed that the 991-stimulated glucose uptake was fully dependent on AMPK in both EDL and soleus using skeletal muscle-specific AMPKα1/α2 double KO (m-α1/α2 DKO) mice (**Fig. 5A, B**). Immunoblot analysis validated AMPKα deficiency and profound decreases in phosphorylation of ACC and TBC1D1 in the absence or presence of 991 in both EDL and soleus in the m-α1/α2 DKO mice (**Fig. 5C, D**). We next assessed the effect of 991, MK-8722 (a structural analogue of 991 [24]), and AICAR on γ3-associated AMPK activity in isolated muscle from WT and γ3 KO *ex vivo*. As shown in **Fig 5E**, γ3-associated activity was increased (∼1.5-2-fold) with 991 or MK-8722 and robustly increased (∼3-fold) with AICAR in EDL from WT mice. Despite minimal γ3-associated activity detectable in soleus, the activity was increased ∼2-fold with 991 in WT mice (**Fig. 5F**). As expected, there was no γ3-associated AMPK activity present in skeletal muscle from γ3 KO mice (**Fig. 5E, F**). In contrast to the effect of AICAR, incubation of EDL with 991 or MK-8722 resulted in comparable increases in glucose uptake in WT and γ3 KO mice (**Fig. 5G**). We also observed that 991-stimulated glucose uptake was similar in soleus between WT and γ3 KO mice (**Fig. 5H**). 991 and/or MK-8722 increased phosphorylation of ACC and TBC1D1 in EDL and soleus with no differences in the levels of phosphorylation between WT and γ3 KO mice (**Fig. 5I-L**). Consistent with the *ex vivo* results, oral administration of MK-8722 (10 or 30 mg/kg body weight) resulted in a comparable blood glucose-lowering kinetics *in vivo* between WT and γ3 KO mice (**Fig. 5M** and **Supplementary Fig. 3**). Taken together, we demonstrate that γ3 is dispensable for the stimulation of glucose uptake and disposal in skeletal muscle in response to 991 or MK-8722 (ADaM site targeted compounds).

**Figure 5:**
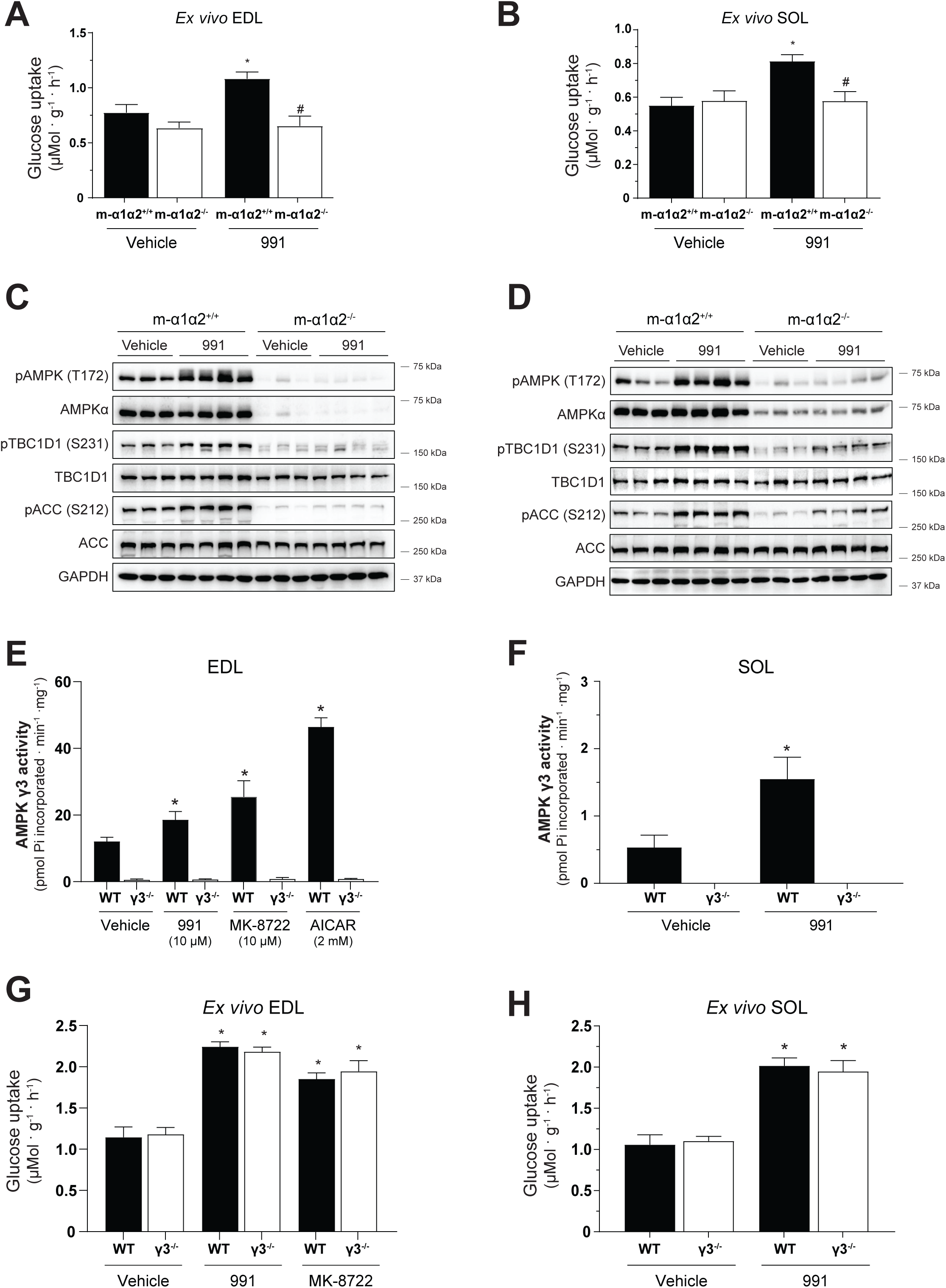
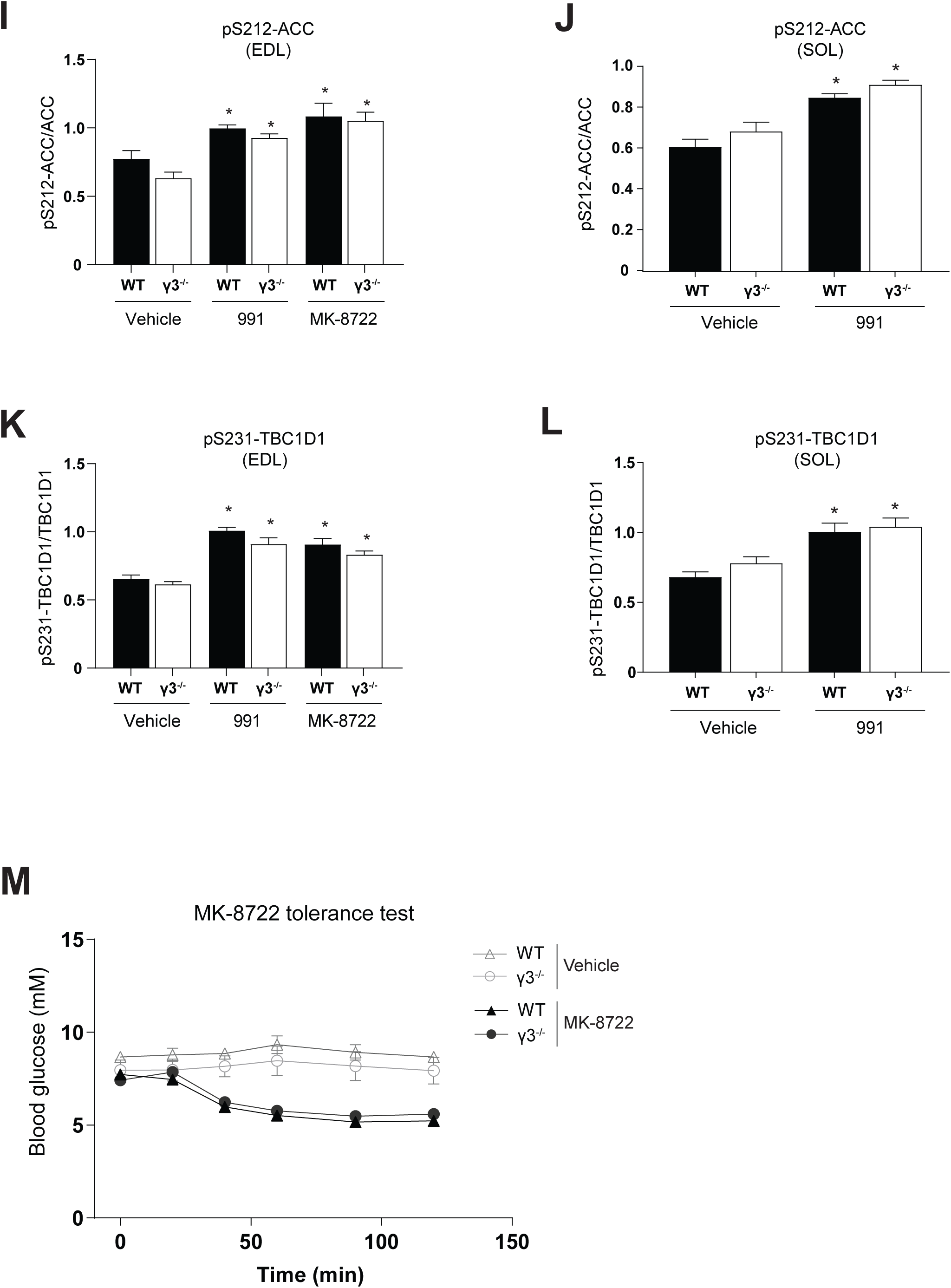
AMPKα1/α2, but not γ3, is required for glucose uptake skeletal muscles and hypoglycemia in response to the ADaM site-targeted activators, 991 and MK-8722. (A-D) EDL or SOL muscles were isolated from the indicated genotypes and incubated in the absence (vehicle, DMSO) or presence of 991 (10 μM) for 50 min followed by an additional 10-min incubation with the radioactive 2-deoxy-glucose tracer. One portion of the muscle extracts was subjected to glucose uptake measurement (A, B) and the other was used for immunoblot analysis using the indicated antibodies (followed by a signal detection using enhanced chemiluminescence) (C, D, n=3-4 per treatment/genotype). (E-L) EDL or SOL muscles were isolated from the indicated genotypes and incubated in the absence (vehicle, DMSO) or presence of the indicated compounds for 50 min followed by an additional 10-min incubation with the radioactive 2-deoxy-glucose tracer. One portion of the muscle extracts was subjected to immunoprecipitation with the γ3 antibody followed by an *in vitro* AMPK activity assay (E, F, n=4-14). The other portion was subjected to glucose uptake measurement (G, H, n=4-9) or immunoblot analysis using the automated capillary immunoblotting system with the indicated antibodies (I-L, n=4-9). M) MK-8722 tolerance test. Mice were fasted for 3 hours and orally treated either with vehicle or MK-8722 (10 mg/kg body weight) followed by blood glucose kinetics monitoring over the indicated duration. Results are shown as means ± SEM. Statistical significance was determined using the unpaired/two-tailed Student’s t-test or one-way analysis of variance with Bonferroni correction and are shown as \**P* < 0.05 (treatment effect within the same genotype), ^#^*P* < 0.05 (WT vs. γ3^−/−^ within the same treatment). EDL; extensor digitorum longus; SOL; soleus, AICAR; 5-aminoimidazole-4-carboxamide ribonucleoside

### 3.6. AMPKγ3 protein and its associated AMPK trimeric complexes are present in mouse BAT

*Prakg3* mRNA are expressed in mouse brown adipose precursors [37], however whether γ3 proteins are expressed and exist as part of functional AMPK trimeric complexes in BAT is unknown. We initially performed a comparison of AMPK subunit/isoform protein expression profiles between skeletal muscle (EDL) and BAT from WT mice, which revealed distinct profiles between the two tissues (**Fig. 6A**). Compared to skeletal muscle, BAT expresses relatively higher and lower amounts of α1 and α2, respectively. The total AMPKα content (assessed by a pan-AMPKα antibody) was similar between the tissues. However, the efficacy of the isoform-specific detection of α1 and α2 proteins by this antibody is unknown. We observed a divergent expression pattern of the β isoforms between the tissues. While skeletal muscle predominantly expresses β2, BAT predominantly expresses β1 (**Fig. 6A**). In contrast, γ1 expression is similar between the tissues. We next immunoprecipitated γ3 from BAT (and GAS muscle as control) and performed either γ3-associated AMPK kinase activity assay or immunoblot analysis to identify α and β subunit isoforms interacting with γ3. As shown in **Fig. 6B**, we detected γ3 and its associated AMPK activity in BAT from WT, but not from γ3 KO mice. Interestingly, γ3 preferentially interacts with α2 and β2 (**Fig. 6C**), whereas γ1 interacts with α1/α2 and preferentially with β1 (**Fig. 6D**). We also observed that γ3 deficiency in BAT did not affect abundance of other AMPK subunit isoforms (**Fig. 6E**). Collectively, we provide evidence that AMPKγ3 protein is expressed in BAT and it forms functional complexes by mainly interacting with α2 and β2.

**Figure 6:**
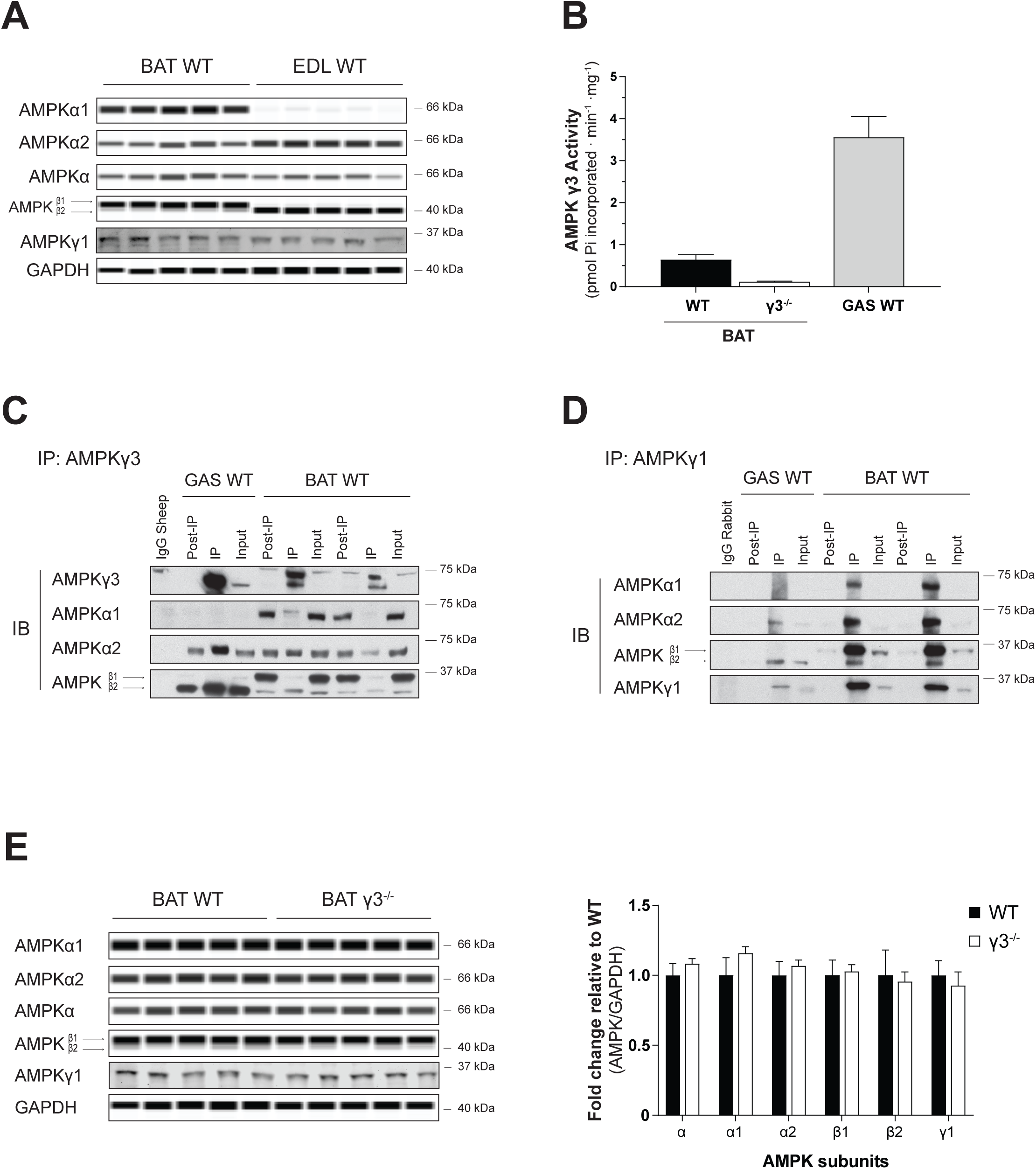
AMPKγ3 is expressed and forms functional trimeric complexes in mouse brown adipose tissue (BAT) (A) Immunoblot analysis of the skeletal muscle (EDL) and BAT extracts harvested from WT mice using the automated capillary immunoblotting system with the indicated antibodies. Note that γ1 expression was quantified using another immunoblotting system (Li-COR) due to antibody compatibility. (B) Extracts from GAS muscle (100 μg) or BAT (1000 μg) were subjected to immunoprecipitation with γ3 antibody and the γ3-containing immune-complexes were assayed for AMPK activity *in vitro*. (C, D) γ3- or γ1-containing AMPK complexes were immunoprecipitated from GAS (100 μg) or BAT (1000 μg) extracts and subsequently subjected to immunoblot analysis using the indicated antibodies followed by a signal detection using enhanced chemiluminescence. (E) Quantification of the isoform-specific AMPK expression of a panel of tissues (harvested from WT or γ3^−/−^ mice) was performed using the automated capillary immunoblotting system with the indicated antibodies. Results are shown as means ± SEM (n=5-7). GAS; gastrocnemius, EDL; extensor digitorum longus

### 3.7. AMPKγ3 is not required for the acute induction of UCP1-mediated non-shivering thermogenesis in the BAT

AMPK plays an important role for BAT formation [37] and thermogenesis in response to cold exposure and β3-adrenoreceptor (β3-AR) stimulation in rodents [36]. We probed BAT function using the β3-AR agonist CL-316,243 (CL), which increases thermogenesis through a UCP1-dependent mechanism [44]. A single injection of CL increased oxygen consumption and interscapular BAT surface area temperature in both WT and γ3 KO mice, but there were no differences between the genotypes (**Fig. 7A-C**). Furthermore, CL increased serum non-esterified free fatty acid concentration to a similar extent in both WT and γ3 KO mice, indicating no major alterations in lipolysis (**Fig. 7D**).

**Figure 7:**
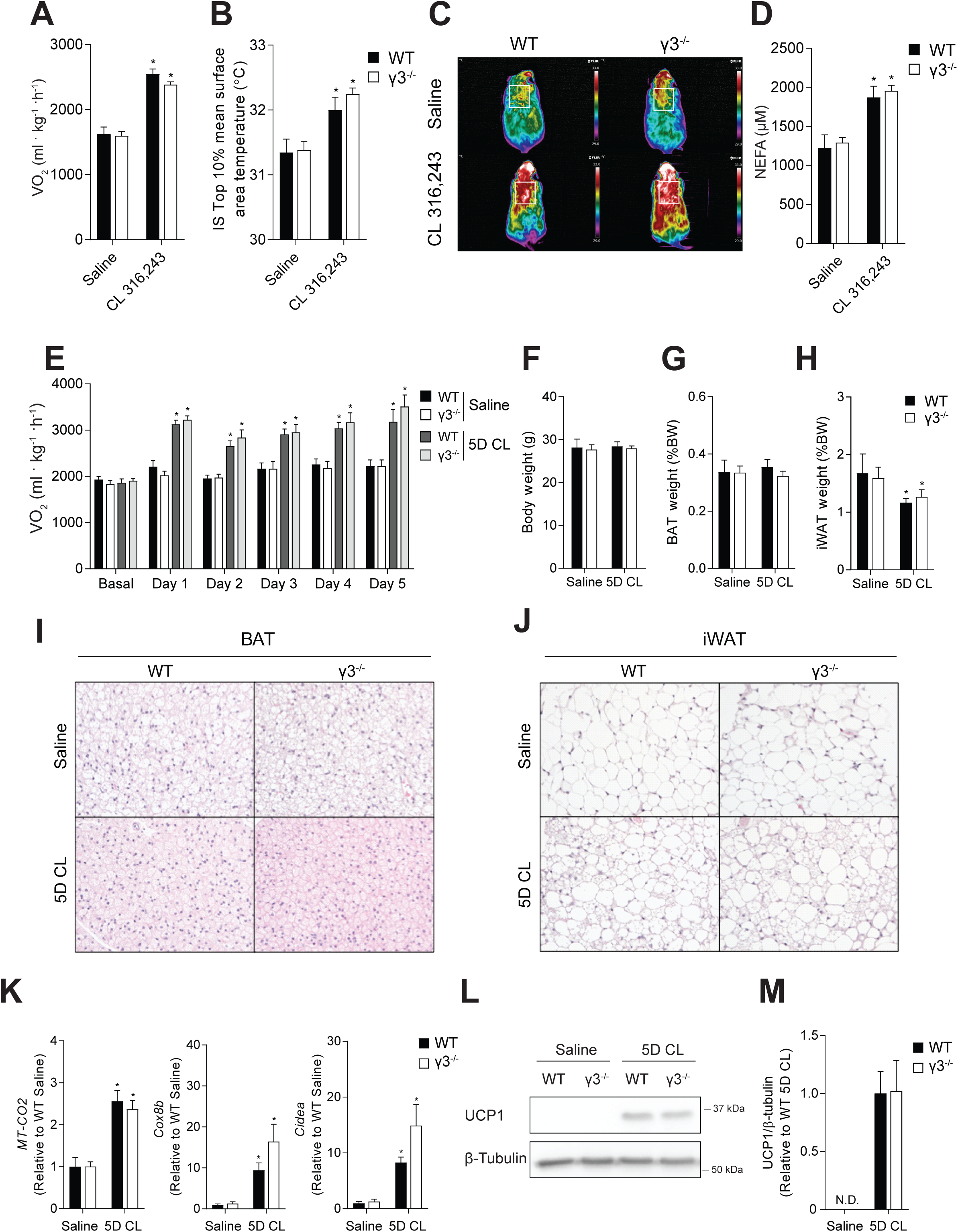
AMPK γ3 is not required for the non-shivering thermogenesis or the browning of inguinal white adipose tissue (WAT) in mice. (A) Oxygen consumption (VO_2_), (B, C) Interscapular brown adipose tissue (BAT) surface area temperature with representative thermal images, and (D) serum non-esterified free fatty acid (NEFA) concentration in response to a single injection of saline or CL 316,243 in male WT or γ3^−/−^ mice (0.033 nmol/g, 20 min time-point), n=9-13 per group. Data are means ± SEM with a CL 316,243 effect shown as \**P* < 0.05, as determined via repeated measures two-way analysis of variance (ANOVA). (E) Oxygen consumption (VO_2_) basally and 6-h post-injection of saline or CL 316,243 in male WT or γ3^−/−^ mice on indicated days, n=5-8 per group. (F) Final body weight (BW), (G) BAT weight, and (H) inguinal WAT (iWAT) depot weight following 5 consecutive days of saline or CL 316,243 (5D CL) injections in male WT or γ3^−/−^ mice, n=5-8 per group. (I, J) Representative histological images of H&E-stained BAT (I) and iWAT (J) (10X magnification) from male WT or γ3^−/−^ mice treated with saline or 5D CL. K) mRNA expression of genes indicative of iWAT browning, *MT-CO2* (n=4-8 per group), *Cox8b* (n=5-8 per group), and *Cidea* (n=5-7 per group) in male WT or γ3^−/−^ mice treated with saline or 5D CL for 5 days. L) Immunoblot analysis and densitometry quantification (M) of UCP1 in male WT and γ3^−/−^ mice treated with saline or 5D CL for 5 days (n=6-8 per group). Data are means ± SEM with \**P* < 0.05 denoting a 5D CL effect, as determined via repeated measures two-way ANOVA (A) and regular two-way ANOVA.

### 3.8. AMPKγ3 is not required for the adaptive response to non-shivering thermogenesis or the browning of inguinal white adipose tissue (iWAT)

We next performed injections of CL for 5 consecutive days to determine if γ3 is required for the adaptive response to non-shivering thermogenesis or the browning of iWAT. Daily treatment of mice with CL increased oxygen consumption without altering body or BAT weight, but did reduce iWAT weight similarly in both WT and γ3 KO mice (**Fig. 7E-H**). Furthermore, γ3 KO mice treated with CL for 5 days had similar morphological changes in BAT – with smaller lipid droplets – and the appearance of multilocular adipocytes within iWAT (**Fig. 7I, J**). Lastly, CL treatment increased UCP1 expression (at both transcript and protein levels), as well as levels of other thermogenic and mitochondrial genes such as *Cox2, Cox8b and Cidea* in both WT and γ3 KO mice (**Fig. 7K-M**). These results demonstrate that γ3 is dispensable for β-adrenergic-induced remodeling of BAT and iWAT in mice.

## 4. DISCUSSION

AMPK has been considered a promising target for the treatment of the metabolic syndrome over the last decades. The identification of new mechanisms for drug-targeting on AMPK (i.e. discovery of ADaM site) has advanced the development of more potent and selective AMPK activators with improved bioavailability [9]. Recent proof of concept studies in rodents and non-human primates have compellingly demonstrated that oral administration of pan AMPK activators (e.g. MK-8722, PF-739) targeting the ADaM site can promote glucose uptake in skeletal muscle, and ameliorate insulin resistance and reduce hyperglycemia without causing hypoglycemia [23; 24]. Since γ3 is exclusively expressed in skeletal muscle, understanding of the physiological roles that γ3 plays in regulating glucose metabolism/homeostasis is important for the development of skeletal muscle-selective AMPK activators. In addition, a recent *in vitro* study reports that γ3 plays a role in BAT development [37] prompted us to investigate the role for γ3 in thermogenesis and adipose browning *in vivo*. In the current study we found that genetic ablation of γ3 resulted in a selective loss of AICAR-, but not MK-8722-induced blood glucose-lowering *in vivo* and glucose uptake specifically in glycolytic muscles *ex vivo*. We also found that γ3 is dispensable for the acute induction of UCP1-mediated non-shivering thermogenesis in BAT or the adaptive response to non-shivering thermogenesis and the browning of WAT.

We observed that the levels of α2 and β2 isoforms were reduced (∼20-30%) in glycolytic (GAS and EDL), but not in oxidative (soleus) muscle in γ3^−/−^ KO compared to WT mice. In soleus, a previous study reported that γ3 was only detectable in a complex with α2 and β2, but relative amount of this complex was shown to be only 2% and >90% of α2 and β2 forms complexes with γ1 [21; 34]. In contrast, α2 and β2 form complexes with γ1 (70%) and γ3 (20%), respectively in EDL muscle. Therefore, it is plausible that a constitutive deficiency of γ3 resulted in a partial loss of α2 and β2 proteins due to degradation of excess monomeric forms of α2 and β2 in GAS/EDL, but not in soleus muscle. Consistent with this notion, a constitutive deletion of γ1, a ubiquitously expressed γ isoform across tissues, resulted in much more profound reductions of all its interacting AMPK isoforms (α1, α2, β1 and β2) in mouse tissues including skeletal muscle [51]. Even though previous work reported that γ3 deficiency did not affect the levels of other AMPK isoforms in GAS muscles, there was an ∼25% reduction of α2 protein expression in GAS muscles from γ3 KO compared to WT mice [25]. It might be the case that it did not reach statistical significance due to insufficient power. In line with this assumption, phosphorylation of AMPKα (Thr172) was reduced in EDL from the same γ3 KO mouse model (compared to WT) [52].

A complete loss of functional α2 or β2 was associated with ablated glucose uptake with AICAR in mouse skeletal muscle *ex vivo* [28; 30; 31]. To our knowledge, whether a partial loss of α2 and/or β2 (e.g. in heterozygous α2^+/−^ or β2^+/−^ mice) reduces glucose uptake in EDL with AICAR *ex vivo* is unknown. However, we have previously demonstrated that a profound reduction (>60%) of α2 activity observed in EDL of the LKB1 hypomorphic mice resulted in comparable AICAR-stimulated glucose uptake and ACC phosphorylation compared to WT mice [4]. Moreover, here we report that 991/MK-8722 stimulates glucose uptake in EDL muscle from the γ3 KO mice. Therefore, a partial reduction of α2/β2 expression (∼20-30%) is unlikely to be responsible for the decrease in AICAR-stimulated glucose uptake in γ3-deficient EDL muscle.

One of the major findings of this study was that γ3-deficiency caused blunted glucose uptake in EDL *ex vivo* and hypoglycemic response *in vivo* with AICAR, but not with ADaM site-targeted compounds (i.e. 991, MK-8722). This was particularly intriguing as both AICAR and 991/MK-8722 require intact AMPK catalytic activity to promote glucose uptake in skeletal muscle tissues/cells [23; 24; 28-32; 45; 50], and we report that both AICAR and 991/MK-8722 increased γ3-associated AMPK activity. The dose of AICAR (2mM) utilized had a more potent effects on γ3-associated activity (∼3-fold increase) as compared to 10 μM 991/MK-8722 (∼1.5-2-fold). However, the results from this assay do not reflect cellular activity, as the *in vitro* kinase assay following immunoprecipitation accounts for covalently-regulated activity (e.g. phosphorylation), but not allosterically-regulated (i.e. by AMP/ZMP, 991/MK-8722) activity. Judging from phosphorylation levels of ACC and TBC1D1, AICAR and 991/MK-8722 comparably increased cellular AMPK activity in skeletal muscle. However notably, compound-induced phosphorylation of ACC and TBC1D1 *ex vivo* was only reduced in EDL when treated with AICAR, but not with 991/MK-8722, in γ3 KO compared to WT. This raises the possibility that AICAR preferentially activates γ3- over γ1-containing complex(es). Concordantly, AMP appears to have stronger binding affinity to nucleotide binding site 3 (in the CBS domain) of γ3 (∼40 μM) than γ1 (∼300-600 μM) *in vitro* (using bacterial AMPK-complex preparations) [53]. On the other hand, another *in vitro* study reported that while AMP potently (allosterically) activated α2β2γ1 (∼3-fold), it barely activated α2β2γ3 (<15%) complex [54]. Thus, how these *in vitro* results can be interpreted and translated into cellular context is unclear. The γ-isoforms all contain a highly conserved C-terminal region harboring the four CBS domains. Conversely, the γ2 and γ3 isoforms contain long N-terminal extensions that are not present in the γ1 isoform. These N-terminal extensions display no apparent sequence conservation between isoforms. To date, there are no crystal structures available for γ2- or γ3-containing AMPK trimeric complexes and it is unknown whether the N-terminal extensions of γ2 or γ3 play any functional role. A recent study using cell-based assays demonstrated that α2β2γ1 and α2β2γ3 complexes were similarly activated in response to 991 treatment, whereas α2β2γ2 complexes exhibited a greater activation (compared to α2β2γ1/α2β2γ3 complexes) [55]. The authors proposed that the effect is mediated by the N-terminal region of γ2 and is due to enhanced protection of AMPKα Thr172 from dephosphorylation. Whether N-terminal extension of γ3 has any specific role to play in muscle cells/tissue and in AMP/ZMP-mediated regulation of AMPK and glucose uptake in skeletal muscle is unknown.

Even though γ3 deficiency was associated with reduced AICAR-stimulated glucose uptake in EDL muscle *ex vivo*, we provide the first evidence that the AICAR-induced blood glucose lowering effect *in vivo* was robustly reduced in γ3 KO compared to WT mice, which was quite similar to AMPKα2 KO and α2 kinase-dead (KD) expressing transgenic mice [28; 30]. Indeed, α2 KO and KD mice still showed decreases in blood glucose in response to an acute injection of AICAR, which is most likely due to the inhibitory effect of AICAR on hepatic glucose production through ZMP-dependent inhibition of fructose 1,6-bisphosphatase 1 [13; 56]. MK-8722 has been shown to cause hypoglycemic effect through the stimulation of glucose uptake in both glycolytic (GAS) and oxidative (soleus) muscle *in vivo* [24]. In contrast to AICAR, but consistent with *ex vivo* data, we provided evidence that MK-8722-induced skeletal muscle glucose uptake and blood glucose-lowering effects were comparable between γ3 KO and WT mice. This compellingly demonstrates that γ3 is dispensable (in other words γ1 is sufficient) in stimulating glucose uptake in skeletal muscle in response to pan-AMPK ADaM site-binding activators.

Evidence suggests that AMPK plays a vital role in regulating the development of BAT, maintenance of BAT mitochondrial function, and browning of WAT [35]. We provided genetic evidence that mice lacking functional AMPK specifically in adipocytes, through an inducible deletion of β1 and β2, were intolerant to cold and resistant to β-adrenergic stimulation of brown and beige adipose tissues [36]. Similar findings were also observed in AMPK α1/α2 KO mice [57]. Detailed protein expression profiles of AMPK subunit isoforms in BAT in comparison to other tissues have not been performed, and to our knowledge presence of γ3 protein in BAT has not been demonstrated. RNA sequencing results identified γ3 at intermediate amounts (Reads Per Kilobase of transcript per Million mapped reads (RPKM), >40) in mouse brown preadipocytes [37]. Strikingly, RNAi-mediated knockdown of either γ1 or γ3 (but not γ2) in brown adipocyte precursors was sufficient to profoundly reduce (>80%) UCP1 protein expression. Using γ3-specific antibodies we developed, we have demonstrated that γ3 protein/activity is present and in complex mainly with α2 and β2 in mouse BAT. Even though we showed that γ3 is dispensable for β-adrenergic-induced thermogenesis and remodeling of BAT and iWAT, future studies are warranted to determine if the specific activation of the γ3-containing complexes (when such drugs are available) induces adipose browning and subsequent amelioration of insulin resistance and fatty liver disease.

## 5. CONCLUSIONS

We demonstrated that a genetic loss of γ3 resulted in a selective loss of AICAR-stimulated glucose-lowering *in vivo* and glucose uptake specifically in glycolytic skeletal muscles *ex vivo*. We also showed that γ3 is dispensable for thermogenesis and the browning of WAT. The potent pan-AMPK activators targeting the ADaM site are effective in reversing hyperglycemia in rodents and non-human primates, and this is due to activation of AMPK in skeletal muscle, not liver [23; 24]. This might make them valuable adjuncts to metformin, which acts primarily on the liver [13; 58; 59]. However, there are remaining important safety issues that need to be carefully considered and examined, such as the potential for AMPK activation to promote cardiac hypertrophy or the survival of cancer cells (e.g. under hypoxic conditions). To avoid these potential liabilities, the development of AMPK activators that can be targeted to specific tissues (for example, liver, muscle and adipose) by taking advantage of isoform-specific selectivity may be beneficial. We and others have shown that selective targeting of specific AMPK isoforms (e.g. α1, β1) by small molecules is possible [9; 60; 61]. When a γ3-complex selective activator is available in the future, ascertaining whether it sufficiently promotes skeletal muscle glucose uptake without causing cardiac hypertrophy and glycogen accumulation will be of interest.

## AUTHOR CONTRIBUTIONS

Conceptualization: K.S. Experimental design: P.Rh., E.M.D., P.R., D.A., N.B., J.S., M.D.S., A.J.O., M.F.K., G.R.S., K.S. Experimental execution: P.Rh., E.M.D., P.R., D.A., N.B., J.S., M.D.S., A.J.O., J.M.Y., A.M.E., J.L.S.G., Q.O., M.F.K., M.M. Supervision: J.S., N.J., J.S.O., J.T.T., P.M., J.W.S., M.J.S, P.D., S.C., G.R.S., K.S. Writing- Original draft preparation: K.S., P.Rh. Writing- Reviewing and Editing: All authors.

## ACKNOWLEDGMENTS

We thank Carles Canto, Magali Joffraud, Guillaume Jacot, Maria Deak, Caterina Collodet, Alix Zollinger, Sylviane Metairon, and Stefan Christen (all affiliated with Nestlé Research) for their technical assistance and input for experimental design, assays, and data analysis/interpretation. We also thank Juleen Zierath for her critical review of the manuscript.

---

**Supplementary Figure 1:**
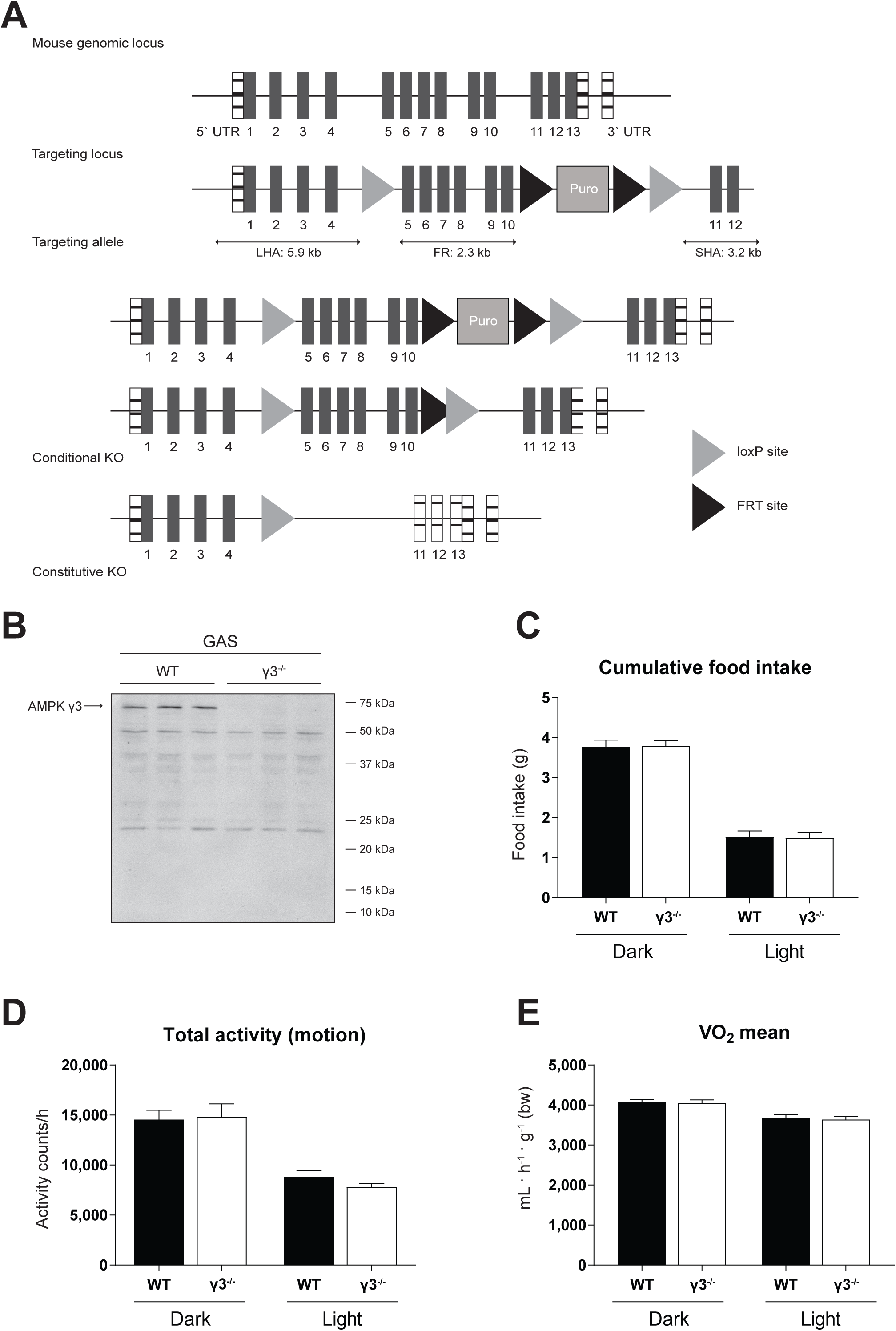
Generation and general characterization of the AMPKγ3 knockout (KO) mice. (A) A schematic illustrating the targeting strategy used to generate Prkag3 knockout (AMPKγ3 KO) mouse model (C57BL/6 background). The targeting strategy is based on NCBI transcript NM_153744_3. The constitutive KO allele is obtained after *in-vivo* Cre-mediated recombination using Cre-Deleter mice (Taconic Biosciences) in which Cre is expressed under the control of the Gt(ROSA)26Sor gene. Deletion of exons 5-10 should result in the loss of function of the Prkag3 gene by deleting the cystathionine β-synthase (CBS) 2 domain and parts of the CBS 1 and 3 domains and by generating a frame shift from exon 4 to exon 11 (premature stop codon in exon 12). In addition, the resulting transcript may be a target for Non-sense Mediated RNA Decay and may, therefore, not be expressed at significant level. (B) Immunoblot analysis of the gastrocnemius (GAS) muscle extracts obtained from the indicated genotypes using the anti-γ3 antibody raised against residues 44–64 (within exon 1-3) of the mouse γ3. (n=3 per genotype) (C-E) General mouse phenotyping was performed by PHENOMIN (Illkirch, France). Mice were housed in metabolic phenotyping cages (TSE system, Labmaster, Germany) and after a 3-hour acclimatization period at ambient temperature (21°C ± 2), food intake (C), ambulatory activity (D) and oxygen consumption (E) were monitored. (n=10 per genotype) Results are shown as means ± SEM.

**Supplementary Figure 2:**
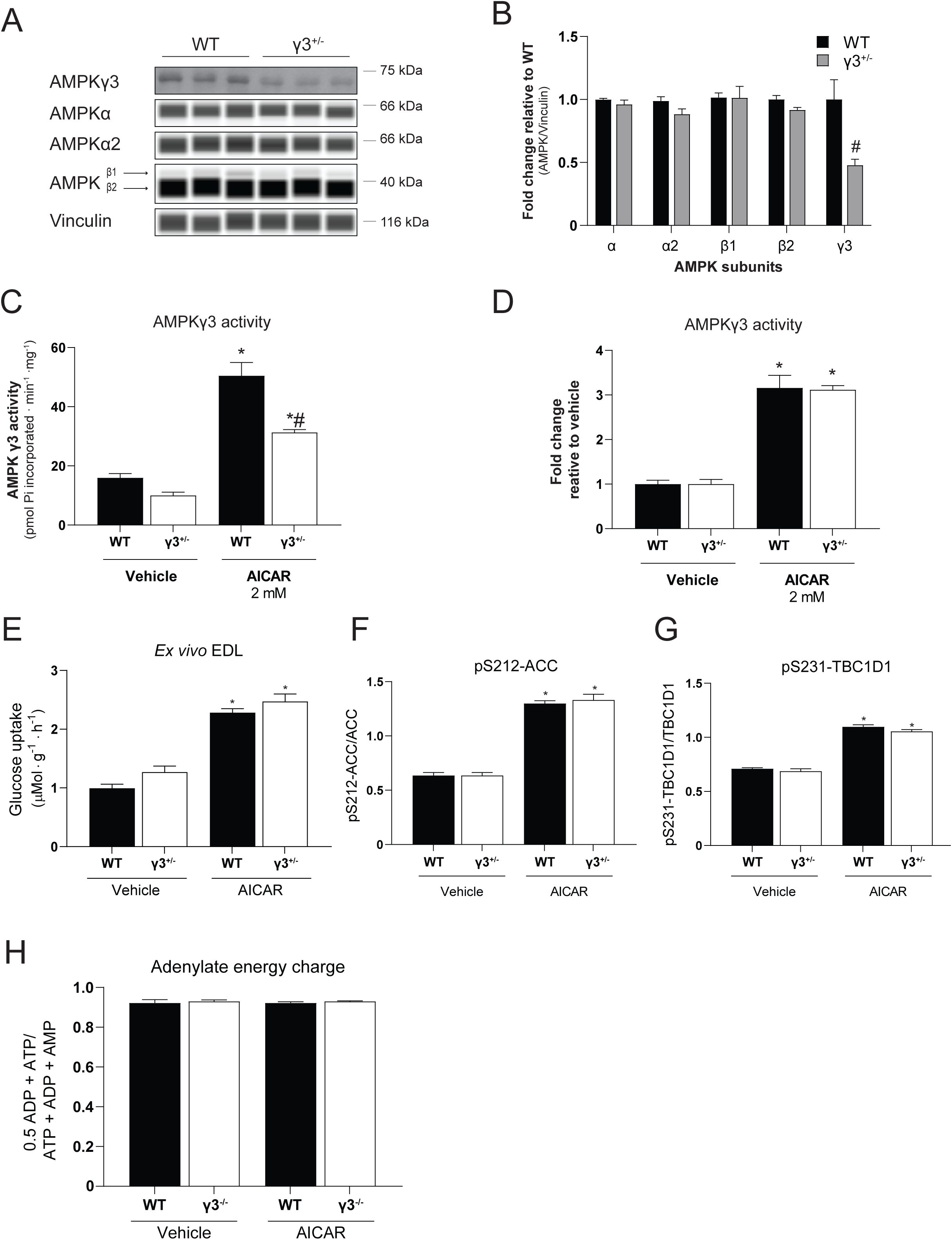
AMPK isoform expression, γ3 activity, AICAR-stimulated glucose uptake and AMPK signaling in heterozygous γ3^+/−^ mice and muscle energy charge following AICAR injection in WT and γ3^−/−^ mice. (A, B) Immunoblot analysis and quantification of the AMPK isoform expression in extensor digitorum longus (EDL) muscles from wild-type (WT) and heterozygous γ3^+/−^ mice using the indicated antibodies. Representative blot images shown (n=5-6 per genotype). (C, D) γ3-containing complexes were immunoprecipitated and were subjected to an *in vitro* AMPK assay. The activity was shown as absolute unit (C) or fold increase relative to vehicle for corresponding genotype (D). (n=5-6 per treatment/genotype) (E-G) EDL muscles were isolated from the indicated genotypes and incubated in the absence (vehicle, DMSO) or presence of AICAR (2 mM) for 50 min followed by an additional 10-min incubation with the radioactive 2-deoxy-glucose tracer. One portion of the muscle extracts was subjected to glucose uptake measurement (E) and the other was used for immunoblot analysis using the automated capillary immunoblotting system with the indicated antibodies (F. G) (n=5-7 per treatment/genotype). H) Following the AICAR tolerance test (Fig. 4G), mice were euthanized and GAS muscles were extracted and nucleotide levels were determined for calculation of the energy charge. (n=5-12 per treatment/genotype. Results are shown as means ± SEM. Statistical significance was determined using the unpaired/two-tailed Student’s t-test or one-way analysis of variance with Bonferroni correction and are shown as \**P* < 0.05 (treatment effect within the same genotype), ^#^*P* < 0.05 (WT vs. γ3^+/−^ within the same treatment).

**Supplementary Figure 3:**
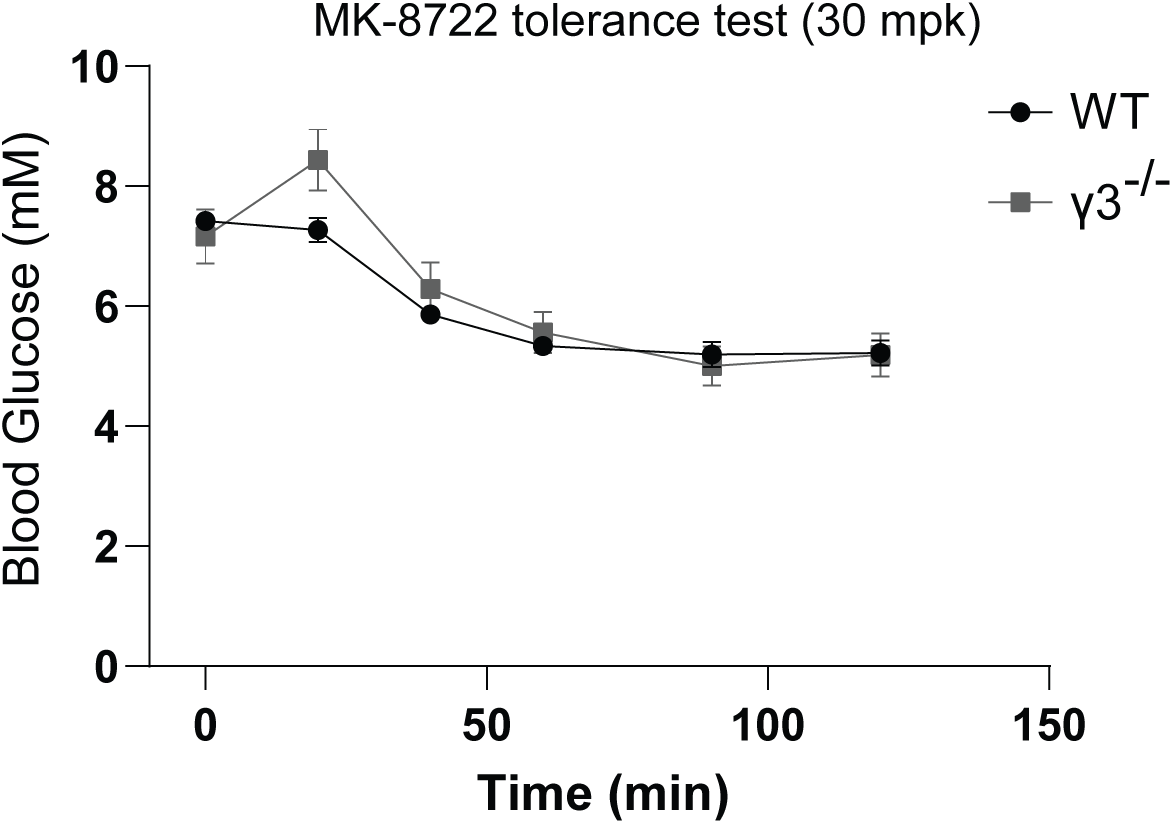
MK-8722 tolerance test in WT and γ3^+/−^ mice. Mice were fasted for 3 hours and orally treated with MK-8722 (30 mg/kg body weight) followed by blood glucose kinetics monitoring over the indicated duration.(n=8-10 per treatment/genotype) Results are shown as means ± SEM.

