## Supplementary Tables for "Compound- and fiber type-selective requirement of AMPKγ3 for insulin-independent glucose uptake in skeletal muscle"

**Supplementary Table 1: list of the primary antibodies**

| Antibody | Application (IP/IB/IF) | Supplier | Reference |
|---|---|---|---|
| AMPKα1 | IP | Custom-made (MRC PPU Reagents & Services, University of Dundee) | PMID: 27577855 |
| AMPKα2 | IP | Custom-made (MRC PPU Reagents & Services) | PMID: 27577855 |
| AMPKγ3 | IP | Custom-made (MRC PPU Reagents & Services) | S673D, PMID: 27577855 |
| AMPKα1/α2 | IB | Cell Signaling | 2532 |
| AMPKβ1/β2 | IB | Cell Signaling | 4150 |
| TBC1D1 | IB | Cell Signaling | 6929 |
| Akt | IB | Cell Signaling | 4691 |
| phospho-Akt (Ser473) | IB | Cell Signaling | 4060 |
| GAPDH | IB | Cell Signaling | 2118 |
| Vinculin | IB | Cell Signaling | 13901 |
| Hexokinase II | IB | Cell Signaling | 2867 |
| GLUT4 | IB | Abcam | ab654 |
| AMPKα1 | IB | MilliporeSigma | 07-350 |
| AMPKα2 | IB | MilliporeSigma | 07-363 |
| pospho-TBC1D1 (Ser231) | IB | MilliporeSigma | 07-2268 |
| UCP1 | IB | Alpha Diagnostic International | UCP11-A |
| β-tubulin | IB | Invitrogen | 32-2600 |
| AMPKβ1 | IB | Proteintech | 26907-1-AP |
| AMPKβ2 | IB | Proteintech | 14429-1-AP |
| UGP2 | IB | Proteintech | 10391-1-AP |
| Total OXPHOS Cocktail | IB | Abcam | ab110413 |
| AMPKγ3 | IP/IB | Custom-made (YenZym) | YZ4698 PMID: 27577855 |
| AMPKγ1 | IP/IB | Custom-made (YenZym) | YZ5115 PMID: 27577855 |
| Myh7 (MyHCI) | IF | DHSB | BA-F8 |
| Myh2 (MyHCIIa) | IF | DHSB | SC-71 |
| Myh4 (MyHCIIb) | IF | DHSB | BF-F3 |
| Laminin | IF | Sigma | L9393 |

IB; immunoblot, IP; immunoprecipitation, IF; immunofluorescence

**Supplementary Table 2: list of the secondary antibodies**

| Name | Supplier | Reference |
|---|---|---|
| Peroxidase AffiniPure Goat Anti-Rabbit IgG | Jackson ImmunoResearch | 111-035-144 |
| Donkey anti-Mouse IgG Alexa Fluor 680 | ThermoFischer | A10038 |
| Goat anti-Rabbit IgG Alexa Fluor 680 | ThermoFischer | A21109 |
| Goat anti-Mouse IgG2b Alexa Fluor 350 | ThermoFischer | A-21140 |
| Goat anti-Mouse IgG1 Alexa Fluor 594 | ThermoFischer | A-21125 |
| Goat anti-Mouse IgM Alexa Fluor 647 | ThermoFischer | A-21238 |
| Goat anti-Rabbit IgG Alexa Fluor 488 | ThermoFischer | A-11008 |
| Anti-Mouse IgG HRP-linked | Cell Signaling | 7076 |
| Anti-Rabbit IgG HRP-linked | Cell Signaling | 7074 |

**Supplementary Table 3: List of primer sequences used for qPCR**

| Name | forward (5’ - 3’) | reverse (3’ - 5’) |
|---|---|---|
| Prkaa1 | AGA GGG CCG CAA TAA AAG AT | CTT TCA AGG CTT CGT CAT CG |
| Prkaa2 | TGA AGC GAG CGA CTA TCA AA | CTT CAC AGC CTC ATC GTC AA |
| Prkab1 | TGC TGC AGG TCA TCT TGA AC | TTG TAC CGG TGT GTT GCA CT |
| Prkab2 | CTT CCT GAG CCC AAT CAT GT | CAG CAG CGT GGT GAC ATA CT |
| Prkag1 | CTC CGC CTT ACC TGT AGT GG | AAG TAG TGG GAC CGA TGC TG |
| Prkag3 | GGA AAC AGC TCC TGT CCT GA | CAT CAA AGC GGG AGT AGA GG |
| Hprt | CAG TCC CAG CGT CGT GAT TA | TGG CCT CCC ATC TCC TTC AT |
| GusB | AAC AAC ACA CTG ACC CCT CA | ACC ACA GAT CGA TGC AGT CC |
| Pgk1 | GGG TGG ATG CTC TCA GCA AT | GTT CCT GGT GCC ACA TCT CA |
| mtDNA-16S | CGT CTA TGT GGC AAA ATA GTG AGA A | CCA GCT ATC ACC AAG CTC GTT |
| mtDNA-16S-Probe | (fuorescein)TAG AGG TGA AAA GCC- (MGB-Q500) | |
| mtDNA-ND4 | CAC ATG GCC TCA CAT CAT CAC | GTG GAT CCG TTC GTA GTT GGA |
| mtDNA-ND4-Probe | (fuorescein)CCT ATT CTG CCT AGC AAA- (MGB-Q500) | |
| PMP22 | TTC GTC AGT CCC ACA GTT TTC TC | ACT CGC TAG TCC CAA GGG TCT A |
| PMP22-Probe | (fuorescein)CGG TCG GAG CAT CAG GAC GAG C- (MGB-Q500) | |
| Titin | AAA ACG AGC AGT GAC GTG AGC | TTC AGT CAT GCT GCT AGC GC |
| Titin-Probe | (fuorescein)TGC ACG GAA GCG TCT CGT CTC AGT C- (MGB-Q500) | |

| Name | Tagman primer reference (Invitrogen) |
|---|---|
| Ppia | Mm02342430_g1 |
| MT-CO2 | Mm03294838_g1 (Invitrogen) |
| Cox8b | Mm00432648_m1 (Invitrogen) |
| Cidea | Mm00432554_m1 (Invitrogen) |
